# Decoding of T cell immunodominance patterns by an integrated mechanistic approach

**DOI:** 10.1101/2024.01.10.574975

**Authors:** Miguel Álvaro-Benito, Esam T Abualrous, Holger Lingel, Stefan Meltendorf, Jakob Holzapfel, Paula de Diego Valera, Jana Sticht, Benno Kuropka, Cecilia Clementi, Frank Kuppler, Monika C Brunner-Weinzierl, Christian Freund

## Abstract

CD4^+^ T cell responses to viral infections are driven by immunodominant determinants. Accessing the peptides defining these determinants in humans is essential for advancing our understanding and potentially tuning immune responses to pathogens. State of the art methods identifying CD4^+^ T cell immunodominant epitopes are constrained by limitations in throughput, performance or both. Here, we leverage on the combined use of a reconstituted antigen processing system and of *in silico* prediction tools to query and study CD4^+^ T cell immunodominance of two model SARS-CoV-2 antigens. We applied this combined platform over a DRB1* panel with broad population and functional coverage to gain mechanistic insights beyond single allotypes. This approach delineates a minimalistic peptide pool (59 candidates) featuring a high immunogenic profile (similar response to 10-fold larger pools) through a representative human sample. Analysis of antigen-encoded and processing-related features on IEDB curated immunodominant peptides reveal that distinct antigen processing mechanisms apply for the Nucleocapsid and Spike model antigens. Notably, the First Bind and then cut mechanism was favored for Nucleocapsid-derived epitopes (up to 55 %), whereas the First Cut and then bind predominated for the Spike (80 %). Together, our results highlight differential processing pathways underlying CD4⁺ T cell immunodominance for distinct viral antigens providing a mechanistic foundation to improve epitope prediction algorithms.

**Significance Statement:** Efficient access to immunodominant CD4^+^ T cell epitopes will inform improved peptide pool design for studying or tuning immunity towards any pathogen. Despite advances in the field, *in silico* epitope prediction remains error-prone and experimental efforts are logistically challenging. Epitope prediction tools are mainly focused on the identification of binding motifs while neglecting key processing steps. Considering antigen processing factors and constraints is expected to improve their performance. We describe and apply a platform for the systematic investigation of antigen processing mechanisms and how they impinge on the selection of CD4^+^ T cell immunodominance. Our results validate distinct immunodominant epitope selection pathways for two model antigens at the human population level.

## Introduction

CD4^+^ T cell immunodominance refers to the preferential and recurrent response of T cells to a subset of peptides presented by Major Histocompatibility Molecules/Human Leukocyte Antigens of class II (MHCII/HLAII). Accessing these peptides not only is essential for elucidating the mechanisms of protective immunity but also for guiding the design of effective vaccines and immunotherapies. T cell immunodominance was originally defined upon recall experiments with peptides following immunization of syngeneic mouse strains and enabled the definition of binding motifs(1). However, follow up studies demonstrated that immunodominant epitope selection implies as well antigen-specific features (e.g., position of the epitope in the antigen), processing conditions (e.g., peptide editing by H2-DM or proteases), the combination of allotypes present in the host, and even T cell precursor frequencies (2, 3). Together, all these variables are known to determine the detectable immunodominant responses.

Peptide binding to MHCII molecules is a prerequisite for CD4+ T cell recognition but it is not sufficient to confer immunodominance. Rather, immunodominance is mainly driven by the selective presentation of antigenic peptides by MHCII molecules. *In vitro* and *ex vivo* experiments suggested that immunodominant peptides primarily were selected via the canonical route of antigen presentation(4). However, alternative processing routes have proven a considerable impact in shaping *in vivo* immunodominance(5). Two molecular mechanisms have been proposed to explain epitope selection for MHCII presentation of pathogen-derived immunodominant epitopes: i) the First Bind and then cut in which flexible regions bind to MHCIIs and remain protected from proteolytic degradation(6, 7), and ii) the First Cut then bind, in which partial degradation of the antigen is a requirement for binding(8). Although epitope prediction algorithms have attempted to incorporate such mechanistic constraints predictive performance remains suboptimal (9–11). Thus, the systematic validation of either model is needed to define generalizable rules that that can be applied to epitope prediction pipelines for improved outcomes.

One of the major challenges when assessing immunodominant responses in humans is the genetic and functional diversity of MHCII allotypes, both within individuals and across populations. Studies aiming at identifying peptides recurrently selected by a given allotype in humans have often focused on the kinetic stability of peptide-MHCII complexes. Those studies are typically restricted to specific allotypes and have limited population-wide implications(8, 12). An alternative, more pragmatic framework considers immunodominance at the haplotype level, emphasizing the collective peptide-binding capacity of MHCII allele combinations rather than isolated allotypes(13). In this context, peptides derived from regions enriched in MHCII supertype binding motifs —and exhibiting conserved epitope-associated sequence features— are considered more likely to be targeted by CD4⁺ T cells (14). The concept of MHC supertypes(15) is supported by promiscuous binding of peptides to MHCIIs, and predictions based in these supertypes are estimated to cover up to 50 % of the total immune response towards any antigen.

Here, we used a panel of representative MHCII allotypes to study immunodominance with a particular focus on gaining mechanistic insights, at both, allele and population level. Using the SARS-CoV-2 Nucleocapsid and Spike proteins as model antigens, we combined predictions from state-of-the-art *in silico* epitope identification tools (9–11) with a reconstituted antigen processing system(16, 17). Our integrative approach facilitated the design of a minimalistic peptide-pool with a broad MHCII coverage and capacity to trigger immunodominant responses. Validation in tailored peptide pools for common HLA-backgrounds confirmed these findings. Furthermore, a systematic analysis of relevant features related to the antigens and their processing revealed diverse pathways for the selection of immunodominant epitopes.

## Results

### Establishing a conceptual framework for studying mechanistic insights of patterns of CD4^+^ T cell immunodominant responses in humans

Pathogen-derived peptides triggering immunodominant responses bind with a high kinetic stability to one or various MHCII allotypes (Figure 1a, left). Peptides triggering these responses are selected through either the First Bind then cut(7, 12) or First Cut then bind(8) mechanism (Figure 1a, right). However, experimental evidence for the selection of immunodominant peptides through the abovementioned mechanistic models in humans is restricted to a single allotype, DRB1*01:01. Therefore, to determine whether these models generalize across other MHCII allotypes, we sought to expand our study to a broader set of HLA-DRB1 (18).

**Figure 1.**
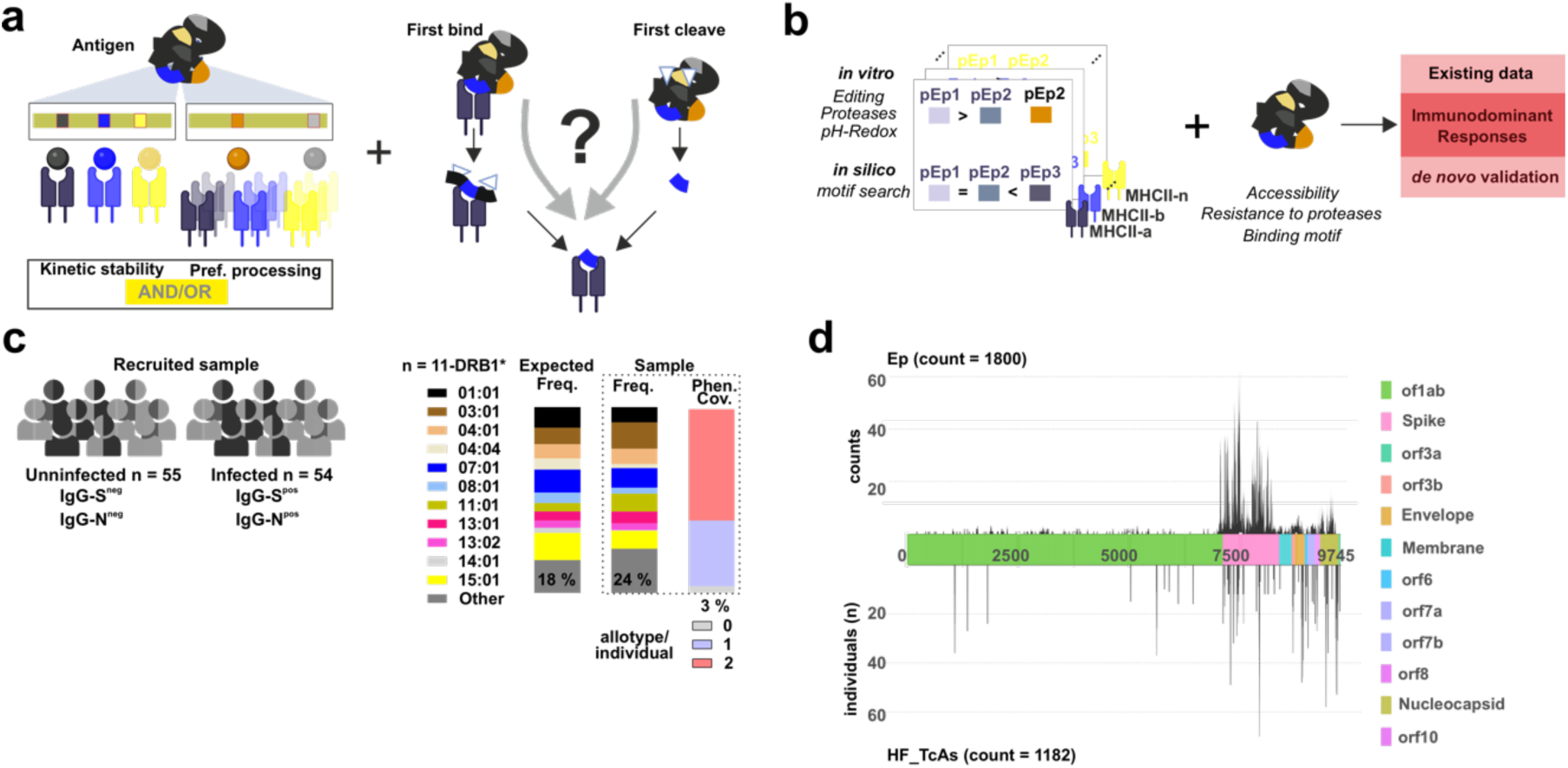
Conceptual and experimental framework to assess key features of immunodominant CD4^+^ T cell responses. **a**. Existing immunodominance frameworks are usually considered mutually exclusive. One of these views is focused on kinetic stability and the other one on promiscuous binding. These views are depicted over a model antigen with 5 potential epitopes (color coded). Depending on the definition of immunodominance considered, each epitope will bind to one allotype (kinetic stability model) or a handful of epitopes will bind to different allotypes (promiscuous binding model). Pathogen-derived immunodominant epitopes are selected through the first bind then cut mechanism or the first cut, then bind. The contribution of each mechanism is not yet very clear (“?”). **b**. We conceived leveraging on predictions from *in silico* tools, identifying binding motifs with the output of a reconstitution antigen processing system resembling the molecular mechanism “first bind then cut” to define immunodominant epitopes. **c**. A sample of 109 individuals whose viral infection status was validated (55 uninfected and 54 SARS-CoV-2 infected) was considered to validate the phenotypic coverage of our DRB1 panel (shown in the right side of the panel) and candidate epitopes. **d**. Overview of reported immune responses to SARS-CoV-2 antigens at the Immune Epitope Data Base (IEDB accessed Sept. 2022, https://iedb.org). The entire viral proteome is depicted in the middle part of the graph indicating amino acid positions and orfs. The count of epitope entries is shown in the upper part of the viral proteome scheme. The number of individuals reacting to peptides defined as High Frequent T cell Assay entries (HF_TcAs, tested in more than 10 individuals and yielding a response frequency higher than 50%) is shown in the lower part of the scheme.

We devised and developed an integrative strategy to study immunodominant epitope selection based on *in silico* tools (assessing mainly binding motifs or peptide presentation) and a reconstitution antigen processing system recapitulating antigen processing (Figure 1b). We selected a panel of 11 DRB1 allotypes providing a phenotypic coverage higher than 90% for Caucasian populations(19) for our 109 donors (96 % phenotypic coverage, Figure 1c, Table S1), validating the use of our panel. The presence of IgG antibodies targeting the Spike protein allowed us classifying individuals as post-Covid19 (PC). Pre-pandemic samples (collected between September 2018 and 2019) were used as un-infected (HD)(20).

To guide our approach, we leveraged existing information of SARS-CoV-2 CD4^+^ T cell responses by retrieving all available datasets from the Immune Epitope Data Base (IEDB) as of September 2022 (Figure 1d upper-part and Figure S1a-e)(21). Relevant studies for our work varied considerably in size (n of individuals tested), broadness (n of antigens tested) and methodology considered for testing immunogenicity. For instance, the high number for T cell Assays (TcAs, n=3437) contrasts with the low count of multimer-identified epitopes (Tet_TcAs, n=149). Entries tested over 10 individuals and reaching Response Frequencies (RF)(22) higher than 0.5, are considered “High Frequency” TcAs (HF_TcAs, Figure 1d lower part). Note that the DRB1* panel reaches a phenotypic coverage on the individuals and datasets collected going from 60% to 80-95 % (Table S2 and S3). Even though most SARS-CoV-2 proteins have proven to be immunogenic we decided to focus on the Nucleocapsid and Spike proteins as main targets of immune responses to viral infection.

### The interplay between *in silico* and *in vitro* predicted epitopes delineates a minimalistic peptide pool with broad population coverage

Most CD4^+^ T cell epitope prediction tools rely on algorithms scoring potential MHCII binders. While these methods offer high throughput, their positive predictive value remains limited. Immunopeptidomics on the other hand implies cellular processing and provides direct information on peptide presentation. However, logistic challenges limit the applicability of this methodology at a broader scale (i.e., sample preparation). We opted for an integrative strategy of *in silico* predictions with experimental methods, allowing us considering antigen processing factors to delineate potential epitopes. We used state of the art *in silico* prediction tools and a reconstituted antigen processing system originally described by Sadegh-Nasseri’s group. This experimental model includes the main steps of the canonical MHCII-antigen processing and presentation pathway(16). We adapted this system to account for the impact of the placeholder peptide Class II Invariant chain peptide (CLIP) on epitope selection and considered leveraging on our quantitative proteomics pipeline(17).

MARIA(9), NeonMHCII(10) and NetMHCIIPan4.0(11), were used to define *in silico* predicted Epitopes (ipEp). Despite their individual utility, these tools showed surprisingly poor concordance in identifying overlapping antigenic regions, with no single tool demonstrating a clear advantage (Figure S2). This observation held true across the entire SARS-CoV-2 proteome. Consequently, we considered the combined output from all three tools (e.g., Figure 2a, upper panel-Nucleocapsid as model antigen for DRB1*03:01, Figure S2a-c). Candidates defined by the reconstituted antigen processing system (experimentally predicted Epitopes, epEp) cluster around specific antigenic regions for each antigen (Figure 2a lower panel-Nucleocapsid as model antigen for DRB1*03:01). We could define MS_H_ and MS_L_ referring to high and low abundance selected candidates (based on MS1 intensities, see Figure S3 and Table S4 for details).

**Figure 2.**
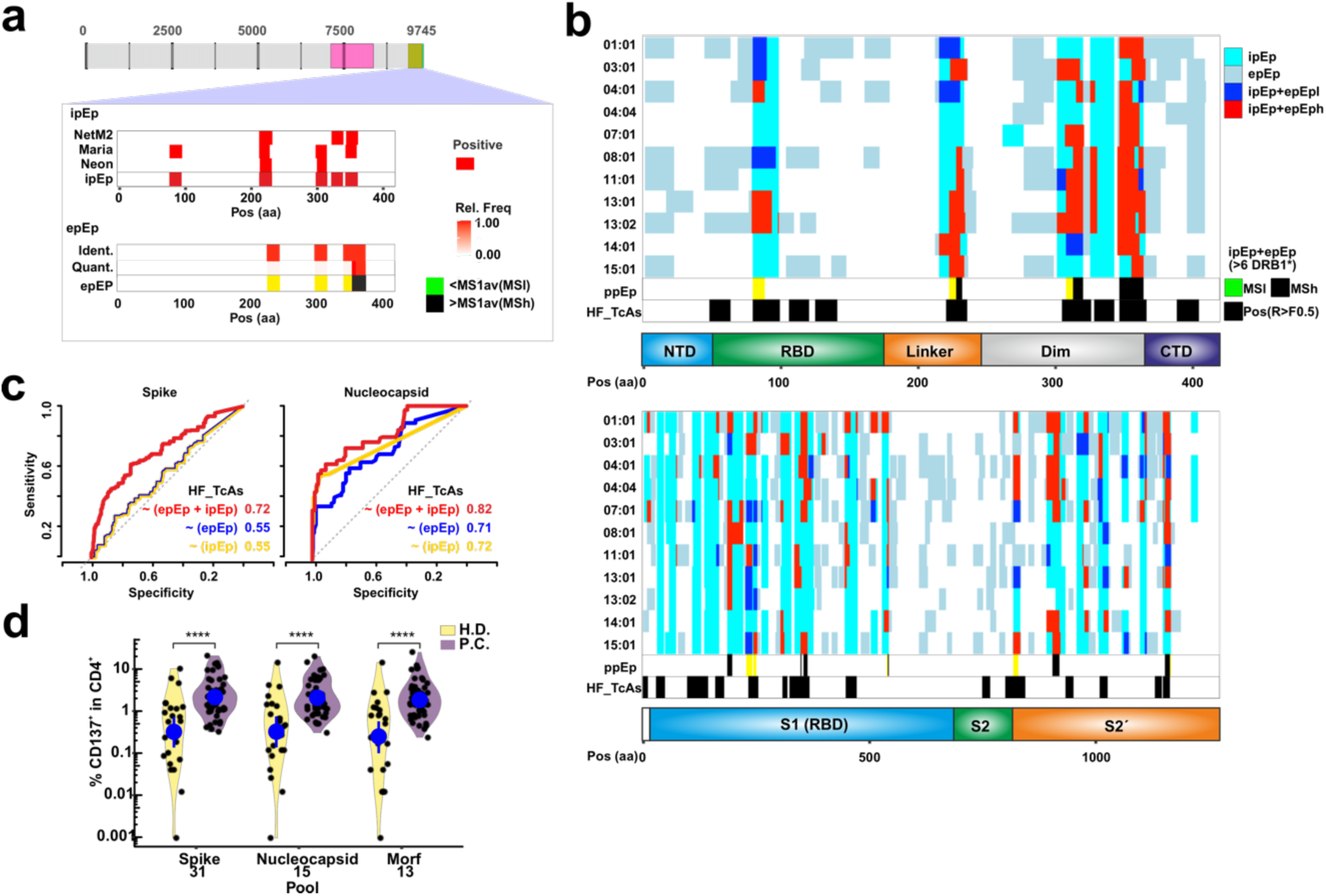
*In silico* and experimental strategies to define potential Epitopes (pEp). **a**. Strategy followed for the identification of potential Epitopes (pEp) by computational and experimental methods. Three *in silico* predictors (NetMHCIIPan, Maria and Neon) are used to determine the likelihood of all potential 15-mers (sliding window of 1 amino acid) from the Spike (pink) and Nucleocapsid (orange) proteins. Candidates identified by any of these tools are considered *in silico* predicted epitopes (ipEp). Experimentally, we applied a reconstituted antigen processing system yielding qualitative and quantitative information and informing regions preferentially selected for presentation, hence experimentally predicted epitopes (epEp). For epEp, if MS1 of a region is higher than the average of the peptides identified by MS then we considered this region as high intensity MSh, and if it was lower MSl. The example shows ipEp and epEp for DRB1*03:01 using the Nucleocapsid protein with each of the *in silico* tools and the outcome of the reconstituted antigen processing system. **b**. Overlap between predictions and experimental data highlights regions of promiscuous predicted Epitopes (ppEp) and existing data on recurrent responses previously measured (HF_TcAs) with a response frequency higher than 50% and tested in at least 10 individuals. Note that we considered here as well MSh and MSl as indicated in a). **c.** ROC curves depicting the performance (as AUC) of ipEp, epEp and their combination to define HF_TcAs from each of the antigens considered based on Binary logistic regression models. See legend for AUC values. **d.** Frequency of activated (CD137^+^) CD4^+^ T cells upon stimulation with each of the pools indicated on the x-axis as determined by flow cytometry. Two groups were tested for each peptide pool, individuals non-previously infected with SARS-CoV-2 (H.D., Healthy donors, yellow, n=24) and individuals that had recovered from SARS-Cov-2 infection (P.C., Post-Covid19, purple, n=48). The difference between the median of the responses was compared applying a non-parametric Mann-Whitney test, and the significance is reported as follows: *p < 0.05; **p < 0.01; ***p < 0.001; ****p < or = 0.0001. Individuals with no response detected where set to 0.001.

Notably, the regions of overlap between epEp and ipEp highlights antigenic regions recurrently selected by several allotypes and aligns relatively well with those classified as HF_TcAs (Figure 2b). Thus, one could assume that the postulated scheme has a decent predictive potential to identify CD4^+^ T cell epitopes. We estimated the probability of classifying residues as immunogenic (HF_TcAs or Tet_TcAs) based on epEp, ipEp and both together, through binary logistic regression models. The Area Under the Curve derived from Receiving Operator Curves for each protein reveals a clear improvement from combining ipEp and epEp (Figure 2c). The low number of Tet_TcAs entries limits considerably the evaluation of the performance of either method for allotype-specific immunodominant epitopes (Figure S4a-b).

Given the benefit of the combination of ipEp and epEp to point out residues contained in immunodominant epitopes, we used their intersection to design a minimalistic peptide pool (MPpN for the Nucleoprotein and MPpS for the Spike) with broad phenotypic coverage. We allowed flexible peptide sizes to increase the likelihood of targeting several restrictions. A subset of peptides from relatively small proteins of SARS-CoV-2, determined exclusively *in silico* (as ipEp), was included as control (Figure S5 and Table S5). Flow cytometry activation assays performed on samples retrieved from our panel of donors revealed a recurrent ability of the three pools to trigger T cell responses in PC individuals compared to HD (Figure 2d). Activated CD4^+^ T cells from PC ranged between 2-15 %, reaching similar frequencies of responders as previously shown by others(23). Increased levels of all cytokines, activation and exhaustion markers tested for all PC individuals when compared to those of HD, validate our minimalistic peptide pools conceived for our broad coverage DRB1-panel (Figure S6).

### Focused validation of immunodominance reveals an alternative mechanistic model of peptide selection

We performed a validation of the immunodominant potential of our MPpN and MPpS candidates to test the performance of our platform, aiming to gain insights into their potential selection mechanisms. To minimize the impact of factors that may bias immune responses (e.g., binding competition), we selected individuals sharing the highest number of DRB1 allotypes, namely DRB1*03:01-*07:01. This combination is present in 2 HD and 3 PC with only partial divergences on HLA-DP (Figure S7a).

Allotype-specific pools feature high and recurrent responses in samples from PC individuals compared to those of HD, as measured via dual-color ELISPOT (Figure S7b-c). The same observations were made using dual peptide combinations. Dual peptide combinations triggering two-fold response frequencies higher than the controls for all PC individuals were considered immunodominant. Two combinations clearly deviate from this criterion (boxed bars, Figure 3a). We defined two subsets of immunodominant peptides: i) those with frequencies of responder cells four-fold higher than any contro for all PC tested individuals (7 combinations, 13 peptides), and ii) those with divergent profiles for every individual (gray and light blue bars respectively, Figure 3a). Most combinations bear at least one peptide with high or medium binding capacity towards either DRB1*03:01 or 07:01 as determined by IC50 measurements (18 peptides out of 23, IC50 < 1000nM, Figure S7d). We assume that peptides with no IC50 value for blocking reporter peptide binding may be presented by any of the additional allotypes from the PC individuals (limited to HLA-DP molecules). Altogether, despite the potential bias in T cell precursor frequencies in these individuals, the observed patterns validate the postulated pipeline and immunodominance of most of the selected candidates.

**Figure 3.**
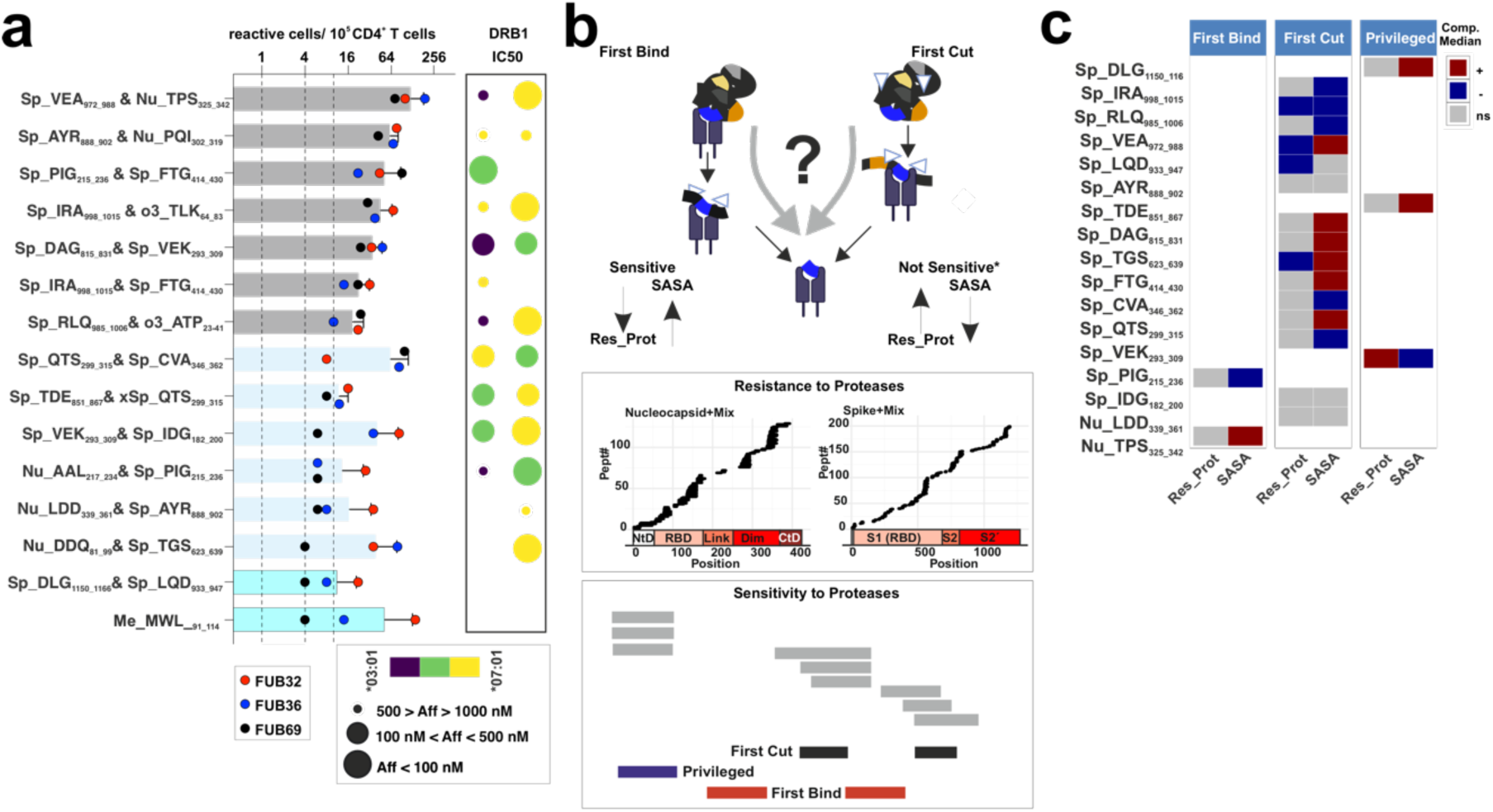
Immunodominance and molecular insights on the selection of allotype-specific peptide pools. **a.** Frequency of responder cells to each dual peptide combination measured via ELISpot (IFNγ+IL-10) from PC donors. The number of spots is converted to responder cells per million of PBMCs and the frequencies determined for every individual (measured in duplicates) are shown according to the color code shown in the legend. Dashed lines represent three minimum thresholds allowing the classification of the measured response. The first line at “10” is the maximum of responder cells detected in any negative control, the second line is the minimum observed response for a combination on the test individuals and the third line represents a 2-fold increase of Line 2. The dual peptide combinations are indicated on the left side and the color code of the bars indicate the distinct immunodominant responses considered (Dark gray and those immunogenic (light gray). Potential restrictions defined by IC50 determination over the two allotypes considered. Each dot represents the affinity of each peptide for either allotype as depicted by the size and color (see legend), Note that affinity differences lower than 1.5-fold are considered as possibly restricted by both allotypes (shown in green). **b.** Antigen-intrinsic and -processing related features defining mechanistic models for peptide selection depicted as scheme. Proteolytic digestion of the two antigens considered reveals peptides resistant to proteases under the tested conditions (Res_Prot) and regions sensitive to proteases that point out at the different mechanistic models. Residues found through more than 3 peptides within series of nested peptides longer than 7 residues are considered indicative of the “First Cut” model. Disruption of series of nested peptides in more than 3 peptides are indicative of the “First Bind” model. Remaining regions with represented in more than 3 peptides with a full coverage of an antigen are considered “Privileged”. **c.** Antigen processing mechanism and antigen-intrinsic features of every peptide tested in the dual combinations from the two model antigens considered. Antigen sources for every peptide are indicated as: Nu-, Sp, o3 and Me for Nucleocapsid, Spike, orf3a and Membrane, respectively. The first three residues of the peptide and the positions are also indicated. Res_Prot and SASA values for each candidate are compared to those of a random selection of peptides excluding all known epitopes (IEDB accession Sept. 2022) and represented according to the scale shown in the right of the panel. “+” indicates higher than median, “-“ refers to values lower than the median and “ns” stands for not significant (significance tested through a Wilcoxon Rank test).

Next, we considered endo-protease cleavage (Sensitivity) and resistance to proteases (Res_Prot, experimentally determined by us, details in Supporting Information and in Figure S8) as well as solvent accessible surface accessibility (SASA) as representative parameters to define the molecular mechanism favoring antigenic peptide selection (Figure 3b). We first evaluated the antigen degradation profiles to determine regions that do require to be protected from proteolysis, hence selected most likely throguh the First Bind model. Then, we delimited those that arise from regions partly or completely resisting proteolysis under the tested conditions and classify them as primarily selected by the First Cut model. Surprisingly, more than 2/3 of the candidates confer to the second mechanism (Figure 3c). A third subset of peptides is identified that could be considered selected via either of the two models since they are capable of withstanding proteolysis under the assayed conditions, and thus we termed them “Privileged”. Altogether, we validate that our MPp have a higher proportion of peptides selected through the First Cut model.

### CD4^+^ Immunodominant epitopes are selected through different pathways for the Nucleocapsid and Spike protein

Our results indicate that the selection of antigenic peptides triggering immunodominant CD4^+^ T cell responses occur primarily through the so called First Cut model, consistent with a previous work focused on Influenzás Hemagglutinin with DRB1*01:01(8). However, the importance of First Bind and then cut mechanism has been recently thoroughly demonstrated(7, 12). Besides, thus far there was no prior report on the so-called “Privileged” peptides that we have described. Under these premises we intended to determine whether and to what extent these observations hold at the human population level for these antigens.

We classified every known HF_TcAs and Tet_TcAs according to the parameter Sensitivity to proteases to validate whether the selection pattern observed for the MPp peptides apply. Surprisingly, we detect a different distribution on the frequency of the models for each antigen. The First Cut model is clearly prominent path for the Spike protein (83 % of the peptides) whereas the First Bind is the preferred one for the Nucleocapsid (55 %) (Figure 4a). Additionally, we observed a very limited frequency of the so-called “Privileged” epitopes (3-4 % for both antigens). Thus, we conclude that the preferred processing pathway for the selection of immunodominant peptides depends on the antigen (Figure 4a lower panel).

**Figure 4.**
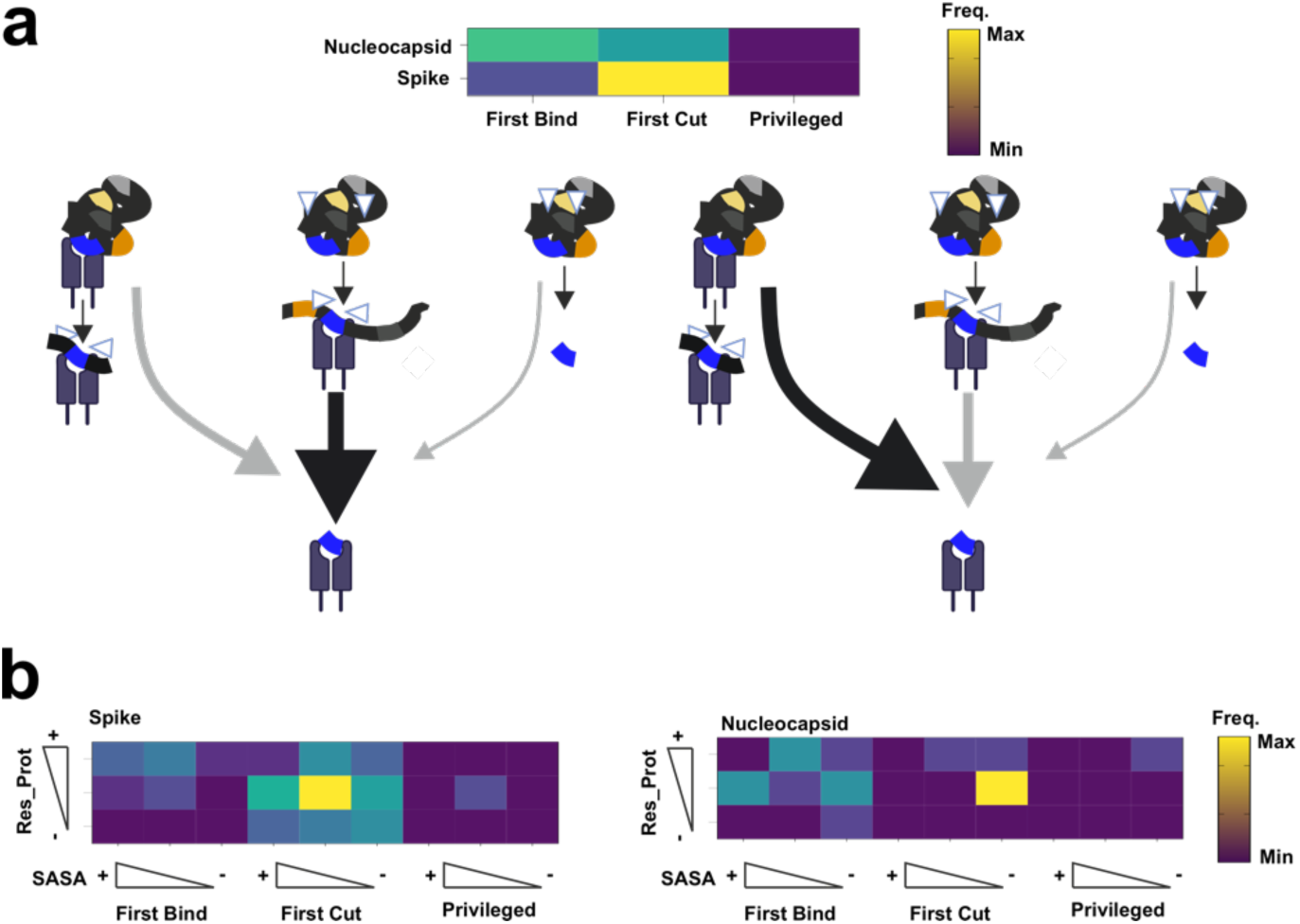
Evaluation of the different mechanistic models through all Nucleocapsid and Spike reported epitopes. **a.** Frequency of peptide selection model for all epitopes previously described at the IEDB (September 2022) for the Nucleocapsid and Spike protein. Each epitope entry for the Nucleocapsid or Spike protein (HF_TcAs or Tet_TcAs) is mapped to the corresponding antigen to define whether they lie at regions sensitive to proteases, as indication of the peptide selection model (“First Bind”, or “First Cut” as well as “Privileged” peptides). Graphical representation of the main findings on the frequency of the peptide selection mechanism for immunodominant epitopes for each antigen. **b**. Heatmaps as in a) depicting the additional impact of SASA and Res_Prot on the entries selected via each molecular mechanism. For each epitope, averaged values for those features were compared to a random distribution of peptides excluding any epitope resulting in three categories. Those categories include higher “+”, no difference “ “ (middle squares), or lower “-“, significance was tested through Wilcoxon-ranked test.

Finally, we decided to evaluate whether any of the additional features considered - SASA and Res_Prot - may inform specific insights for the selection of peptides by each mechanism for each antigen. We accounted for the differences on these features compared to the control peptide pool to conclude that the only apparent difference is that peptides selected through the First Cut mechanism are clearly selected from low SASA regions for the Nucleocapsid protein while the selection from the Spike protein does not follow such pattern (Figure 4b). In conclusion, these results derived from hundreds of human responses indicate that antigen-specific features impose distinct selection patterns.

## Discussion

Our work establishes and validates a robust pipeline for the mechanistic analysis of human CD4^+^ T cell immunodominance. By integrating experimental and *in silico* information across a panel of MHCII allotypes with diverse binding specificities and high population coverage, we provide new insights into the selection and presentation of immunodominant epitopes using the Spike and Nucleocapsid proteins of SARS-CoV-2 as model antigens.

We demonstrate that while *in silico* predictions remain limited by algorithm-specific biases, our experimental workflow effectively narrows down antigenic regions of interest, even across MHCII allotypes with divergent binding grooves. The combination of the two approaches enables the rational design of a minimalistic peptide pool featuring a remarkable population-wide immunodominance profile. Contextualizing antigen and antigen-processing features of peptide candidates confirms that CD4^+^ T cell immunodominance is not restricted to a single molecular mechanism, and that that the specific mechanism depends on the antigen considered. Altogether, we conclude that alternative pathways for epitope selection play a key role and should be considered for improving immunodominant epitope discovery.

Identifying immunodominant CD4^+^ T cell patterns at the human population level has important implications for basic and applied research. Allotype-specific immunodominance has limited applicability at the population level unless broad panels of MHCII allotypes are considered. The use of overlapping peptide pools assuming haplotype restriction capacity on the other hand, lacks insights on restrictions and often neglect binding registers. Our strategy integrates both views of immunodominance not only to identify selection patterns, but also to score how mechanistic insights for pathogen-derived epitopes(7, 8, 12) apply at a population-wide level. Our choice of antigens was motivated by the availability of data, with more than 1800 CD4^+^ epitope entries and approximately double the number of T cell assays (available at the IEDB by September 2022). Besides, our DRB1* panel demonstrates an extraordinary phenotypic coverage throughout our donors but also in the most representative studies published so far (more than 80 % in(23)) enabling the comparison of the extent of the responses detected. We capitalize on the fact that SARS-CoV-2 reactive T cells expanded during the first months of the pandemics are expected to result from naïve T cell clones since no previous contact to this pathogen had been reported although cross-reactivity with other coronavirus antigens is expected (24–29).

We have previously demonstrated the effectiveness of the scheme used here for epitope mapping on complex antigenic mixtures consisting of more than 250 antigens(17). Sadegh-Nasserís group, originally describing the system, has used to map viral epitopes with a mechanistic focus(7, 12, 16, 30). However, this group uses “DRB1* molecules missing the placeholder peptide CLIP, an aspect that can have a considerable impact on epitope selection (reviewed in(31, 32)). This reconstituted antigen processing system, especially when using DRB1* molecules including CLIP, provides direct evidence for peptide selection under conditions resembling cellular processing(6, 12), similar to immunopeptidomics from infected or antigen fed cells(33, 34). In parallel, we employed state-of-the-art epitope prediction tools to capture potential epitopes that might be missed by mass spectrometry or fall outside the specific conditions of our experimental system (e.g., due to the presence of CLIP as placeholder peptide and conditions resembling the first bind model). Surprisingly, our analysis showed a very limited performance for individual prediction tools (Figure 2 and S2). For instance, the prediction outcome of MARIA, deviates considerably from those of NeonMHC(10) and NetMHCIIPan(11), underscoring the lack of consistency across platforms. These discrepancies likely stem from differences in training datasets, algorithmic design, and input parameters—factors that lie beyond the scope of this study (for details see references(9–11)). Despite these limitations, immune response data compiled in the Immune Epitope Database (IEDB) supports the superior performance of our experimental system. Moreover, integrating this experimental output within silico predictions significantly enhances epitope identification, validating the strength of our combined approach.

Comprehensive coverage of the studied antigens using overlapping 15-mer peptides would require roughly 2,000 peptides. In contrast, covering all potential binding registers, or the region estimated to account for 50 % of the total response to these antigens(35) would reduce this number to 550 or 454 peptides, respectively. Of note, the two later strategies overlook specific binding registers or regions essential for optimal T cell recognition(36–38). By prioritizing experimental information and allowing different peptide lengths our approach achieves 10-fold reduction of peptide numbers, and more importantly reached similar T cell activation levels as those reported with the lowest degree of overlap under very similar experimental conditions (measured as % of CD137^+^ in CD4^+^ cells in the range of 0.5-10 %)(39, 40). Thus, our strategy clearly represents an optimization for the assessment of immunogenic regions. Targeted validation of the immunodominant potential of selected peptides was performed on a very specific MHCII background (DRB1*03:01/DRB1*07:01) to prevent MHCII binding competition. The set of 23 peptides (over 59) consisted of two sub-pools covering both DRB1* restrictions present in the samples. These candidates displayed an excellent performance for re-calling immune responses. Potential restrictions were proposed by measuring the ability of each peptide to block reporter binding to each allotype. 6 peptides out of 23 had neglectable or very low binding affinity for the DRB1* allotypes considered under the assayed conditions but they are capable to activate CD4^+^ T cells (S_933-947_, and S_414_430_, S_623_639_, S_1150-1166_, N_339-361_ and M_91-114_). Since most of them had been identified as eluted from these DRB1* molecules in the experimental approach (all except the M_91-114_) we conclude that they represent poor-binders and/or intermediate products of antigen processing hot-spots (e.g. preferentially selected upon binding and proteolysis), or peptides mis-assigned for building the final candidate list. Exemplarily, S_937-951_ appears strongly selected in the DRB1*07:01 experiments (up to 0.17 of the relative MS1 intensity) but our final list includes S_933-947_. Together, these results and observations validate the performance of the proposed approach but highlights that there is still room for improvement. Future studies should aim at scoring the tradeoffs of different peptide lengths for covering broader spectrum of MHC backgrounds that may facilitate clinical applications related to diagnostics or prevention.

Antigen processing constraints play a pivotal role in peptide selection by MHCIIs, thereby facilitating epitope discovery(7, 30, 41, 42). Our analysis confirms that known epitopes and those that we predict exhibit features previously associated with different molecular mechanisms (Bind. Motif, Res_Prot and SASA). Two recent studies have demonstrated the impact of antigen processing constraints on the selection of immunodominant epitopes from Infuenza’s Hemagglutinin(8) and almost the entire HIV proteome(12). Cassotta *et al.*(8) used an Antigen Processing Likelihood(41) estimate to conclude that immunodominant peptides are predominantly located close to unstable protein regions. Proteolytic cleavage at endosomal compartments is required for binding implying selection through a mechanism that could be interpreted as First Cut then bind. Sengupta and collaborators(12) on the other hand report that immunodominant peptides lie mainly at unstable regions that can bind directly to MHCIIs. Additionally, low processing and competition for biding result in poor selection of conserved immunodominant epitopes. Previously, the same group reported that pathogen derived epitopes would be relatively sensitive to proteases(7). Besides querying the presence of binding motifs and solvent accessible surface area (SASA)(41, 42), we determined experimentally proteolytic degradation. This additional parameter is considered key for immunodominant epitope selection. Interestingly, we followed essentially the same experimental approach as Sengupta and collaborators albeit our results point at different molecular mechanism for the selection of immunodominant peptides. Thus, we reason that the presence of CLIP as placeholder peptide in our scheme impose the selection of specific peptide repertoires. Future works expanding our knowledge on CD4^+^ T cell epitopes(21) combined with tools as AlphaFold enabling efficient structure predictions(43) will probably equip us with more accurate information.

We demonstrate the potential of integrating experimental and *in silico* approaches to effectively assess and understand immunogenicity. Our findings underscore that the impact of antigen-related features (e.g. SASA) and experimentally definable processing constraints (e.g. proteolytic degradation) should be funneled into the next generation pipelines of effective MHCII-immunodominance prediction. Integrating these factors in predictive models could significantly enhance the precision of epitope identification, ultimately improving our capacity to design vaccines and immunotherapies tailored to individual HLA haplotypes. This integrated perspective may prove particularly valuable for the development of personalized T cell-based responses in the face of future pandemics or in the context of cancer immunotherapy.

## Materials and Methods

### Peptides, viral antigens and recombinant proteins

Peptides and viral antigens were purchased from different vendors. Recombinant proteins were produced as originally described in(44) using a Baculovirus expression system. For details see Supporting Information (SI).

### Antibodies and reagents used in cell culture experiments

All antibodies were commercially acquired, clones and vendors are referenced in the Supporting Information (SI).

### *In silico* and experimental epitope prediction

The SARS-CoV-2 Nucleocapsid and Spike (Uniprot: P0DTC9 and P0DTC2, respectively) were considered as the basis for all our analyses Peptide-MHC class II binding/presentation prediction was performed using three different algorithms, netmhcIIpan(11), maria(9), and neonmhc2(10), with percentile score cut-off of 10 %. Immunogenicity scores used as flagging criteria were determined using the IEDB tool(14) applying an immunogenicity score cutoff value of 50 %, yielding 1643 immunogenic peptides.

The previously described cell-free reconstituted *in vitro* system(16) was modified according to the specific needs of the experiments(17). In all cases our recombinant MHCII molecules include the placeholder peptide CLIP chaperoning their binding groove. Antigen was incubated with recombinant MHCII molecules in the presence of DM and subsequently Cathepsins were added. Immunoprecipitation and peptide isolation was done according to standard procedures. Isolated peptides were processed and subjected to Mass Spectrometry analysis.

### Peptide binding affinity measurements

Peptide binding affinities to HLA-DR molecules were determined in competition experiments. Binding of fluorescently labelled reporter peptides, specific for each allotype tested, was determined in the presence of titrated concentrations of individual un-labelled peptides. Details are provided in the Supplemental Information (SI).

### Proteolytic degradation of antigens

Spike and Nucleocapsid proteins were incubated in the presence of cathepsins under the same conditions as used for the reconstituted antigen processing system but in the absence of MHCII molecules and DM. Samples were divided for their testing in SDS-PAGE, Western blotting and peptide identification by MS.

### MS measurements and data analysis

All samples were initially cleaned up by reverse phase C18 enrichment and measured as previously described. For data analysis we used MaxQuant (v2.0.3.0) with implemented Andromeda peptide search engine. Reconstituted antigen processing system and antigen proteolytic degradation experiments were searched using specific protein databases. A False Discovery Rate of 1% was considered. Identified peptides were processed using PLAtEAU as previously described(17) for the reconstituted antigen processing system, and spectral counts were considered for the proteolytic degradation. More details can be found in the Supporting Information (SI).

### Donor recruitment, HLA-typing and determination of the infection status

Evaluation and approval of compliance with regulations was requested to, and approved by the Ethics committee of the Universitätsklinikum of the Otto-von-Guericke Universität Magdeburg (UOvG). Donors were recruited at the Medizinische Fakultät/Universitätsklinikum of UovG. Informed consent was signed and agreed upon according to the protocol approved. HLA-typing was performed by the DKMS-LSL facility. Samples from healthy individuals had been collected before the SARS-CoV-2 outbreak. SARS-CoV-2 infection status was determined by detection of antibodies targeting the Spike protein. PBMCs were isolated from blood samples of the corresponding donors via density-gradient sedimentation. Isolated PBMCs were either frozen in medium supplemented with 10 % DMSO or further processed for isolation of monocytes and CD4^+^ T cells.

### Reactivity assays: Intracellular Staining and ELISpot

Presence of reactive cells was determined by either cytokine intracellular staining or dual secretion ELISpot. For details see Supplemental Information (SI). Samples in flow cytometry assays were acquired on a FACSCanto II flow cytometer with FACS-Diva software (v10, BD Bioscience). Secreted cytokines in ELISpot assays were determined according to the manufacturer’s instructions and counting of spots was done on a ImmunoSpot analyzer (Cellular Technology).

### Data retrieval and analysis of antigen processing rules and antigenic peptide features

Available entries in the IEDB described as epitopes, ligands or T cell Assay related (TcAs) were coded as binary vectors (0: not present, 1: part of a hit) (Accessed Sept 2022). The sum of appearances of each residue as a Hit facilitates defining their relative frequency. Each candidate epitope identified by the *in silico* predictors or reconstituted antigen processing system were also coded at the amino acid level (details in Supporting Information, SI). Antigen and antigen-processing related features (Sensitivity, Res_Proteases and rel_SASA) were also coded at the residue level. For the context dependent analysis, we determined the average for each parameter according to the values of each amino acid. As control we used a randomized pool of peptides excluding the selected regions.

### Statistical analysis

All analysis were carried out using R, version 4.2.1 over the RStudio suite unless otherwise stated.

## Acknowledgments

The authors are thankful to all other members of the working groups, and to Eliot Morrison for critically reading the manuscript. The authors are also thankful to the Biosupramol Core facility. This project was funded by a grant from the BMBF (01KI2072) to MCBW and CF, TRR186 (project A21N) and FR-1325/20 to CF. M.Á-B acknowledge funding from the DKMS JHRG2022-01, CAM-2022-T1/23572, ISCIII PI24-00605, and a grant of the Fundación Ramón Areces. Funding was also received from BMBF LongCoCID (01EP2101C) (to MCBW) and the Covid19 program by the state of Saxony-Anhalt (FP1) to HL and MCBW.

## Supporting Information

## Materials and methods

### *In silico* candidate epitope prediction

The SARS-CoV-2 genome sequence was obtained from the NCBI database https://www.ncbi.nlm.nih.gov/nuccore/1798174254. We extracted the sequences of the proteins orf1ab, S, orf3a, E, M, orf6, orf7a, orf7Bb, orf8, N, and orf10 based on the reference genome and corresponding to those reference in the main text. We used a sliding window size of fifteen amino acids and a step of one amino acid for the following analysis (9591 peptides). Potential SARS-CoV-2 epitopes were identified using a novel selection workflow based on the integration of prediction algorithms for peptide-MHC class II binding and immunogenicity. The peptide-MHC class II binding/presentation prediction was performed using three different algorithms, netmhcIIpan(1), maria(2), and neonmhc2(3), with percentile score cut-off of 10 %. Immunogenicity scores used as flagging criteria were determined using the IEDB tool(4) applying an immunogenicity score cutoff value of 50 %, yielding 1643 immunogenic peptides. Potential SARS-CoV-2-derived epitopes were identified as the top-ranked overlapping candidates for each allotype of the eleven MHC class II allotypes. Next, allele-specific lists of peptides with a minimum length of 15 residues were defined for each SARS-CoV-2 protein.

### Peptides and viral antigens

Peptides were purchased from GL Biochem (Shanghai) Ltd (10 mg purity > 95 %). Lyophilized peptides were diluted at a final concentration of 10mM in DMSO and subsequently diluted in PBS. When necessary, the pH was adjusted to 7.4. Nucleocapsid and Spike Glycoproteins expressed in HEK cells were purchased from Sino biological (Cat Numbers. 40588-V08B for Nucleocapsid and 40589-V08B1 for Spike) as C-terminal His-tagged. Lyophilized proteins were reconstituted according to the manufacturer specifications.

### Protein methods

MHC proteins (HLA-DRs and HLA-DM) were expressed as previously described(5). Briefly, HLA-DR cDNAs are cloned into the pFastBacDual vector and include Leu-Zippers in their C-termini as well as a sequence encoding for the CLIP peptide followed by Thrombin cleavage site and a G4S linker in the N-termini of the DRB1 polypeptides. Furthermore, the DRA polypeptide encodes a Biotin Acceptor Sequence at the C-termini of the corresponding Leu-Zipper. Expression was achieved by infection of Sf9 cells at an MOI of 5 and harvesting the cells after 4 days. Protein purification was achieved by immunoaffinity chromatography using a L243-FF-Sepharose resin casted in house. In all cases HLA-DR proteins were cleaved with Thrombin and gel-filtrated using a Sephadex S200. For the reconstituted *in vitro* system, these proteins were also cleaved with V8 protease to remove the Leu-Zippers. HLA-DM cDNAs are cloned into pFastBacDual and include a Flag-Tag in the C-termini of HLA-DMA chain. Purification in this case was achieved using an immunoaffinity M2-Sepharose resin, protein was eluted using Glycine pH3.5. After dialysis, the protein was concentrated (Vivaspin MWCO 10kDa) and gel filtrated (Sephadex S200).

### Peptide binding affinity measurements

Peptide binding affinities to selected HLA-DR molecules were determined by competition experiments using fluorescently labelled reporter peptides. Reporter peptide binding signal was measured by FP. HLA-DR molecules expressed in insect cells were thrombin cleaved to facilitate peptide exchange. Competition experiments were set by adding 100 nM HLA-DR, 100 nM reporter peptide (CLIP-FITC for DRB1*07:01 or MBP-FITC for DRB1*03:01) and tittered concentrations of the corresponding peptide in 50mM Citrate Phosphate buffer containing 150mM NaCl at pH 5.3. Each reaction was measured after 12h incubation at 37° C, and the corresponding IC50 values for each peptide were retrieved by fitting a sigmoidal function to the obtained data points.

### *In vitro* reconstituted antigen processing system

The previously described cell-free reconstituted *in vitro* system(6) was modified according to the specific needs of the experiments(7). HLA molecules together with the candidate antigens and the HLA-DM were incubated for 2 h at 37° C in citrate phosphate 50 mM pH 5.2 in the presence of 150 mM NaCl. Cathepsins were added to reaction mixtures after incubation with L-Cysteine (6 mM) and EDTA (5 mM). The final reaction mixture was incubated at 37° C for 2 to 5 hours. Afterwards the pH was adjusted to 7.5, and Iodoacetamide was added (25 mM). Immunoprecipitation (IP) of the pMHCII complexes was performed using L243 covalently linked to Fast Flow sepharose. Peptides were eluted from purified MHCII adding TFA 0.1 % to the samples. Peptides are separated from the MHCII molecules by using Vivaspin filters (10 kDa MWCO). Cathepsin B (Enzo), H (Enzo) at a molar ratio of 1:250, and S (Sigma) at a molar ratio of 1:500, respectively were used in these experiments.

### Proteolytic degradation of antigens

Spike and Nucleocapsid proteins were incubated in the presence of cathepsins in the molar ratios indicated above. Reactions were performed at 37 °C citrate phosphate 50 mM pH 5.2 in the presence of 150 mM NaCl and stopped at t = 0, and t = 3 h, by adding Iodoacetamide, immediate transfer of the samples to ice. The pH was then adjusted to 7.5 by adding Tris-HCl 1 M pH8.0. Samples were splitted, and used for SDS-PAGE analysis, Western blotting and peptide identification by MS. For MS analysis, samples were dried in a SpedVac and treated as described below.

### LC-MS measurements

All samples were initially cleaned up by reverse phase C18 enrichment. The eluates were dried in a SpeedVac, and peptides were reconstituted in 20 ml of H_2_O containing acetonitrile (4 %), and TFA (0.05 %). 6 ml of these mixtures were analyzed using a reverse-phase capillary system (Ultimate 3000 nanoLC) connected to an orbitrap connected to a Q Exactive HF mass spectrometer (Thermo Fisher Scientific). Samples were injected and concentrated on a trap column (PepMap100 C18, 3 μm, 100 Å, 75 μm i.d. × 2 cm, Thermo Fisher Scientific) equilibrated with 0.05 % TFA in water. After switching the trap column inline, LC separation was performed on a capillary column (Acclaim PepMap100 C18, 2 μm, 100 Å, 75 μm i.d. × 25 cm, Thermo Fisher Scientific) at an eluent flow rate of 300 nL/min. Mobile phase A contained 0.1 % formic acid in water, and mobile phase B contained 0.1 % formic acid in 80 % acetonitrile/ 20 % water. The column was pre-equilibrated with 5 % mobile phase B followed by a linear increase of 5–44 % mobile phase B in 70 min. Mass spectra were acquired in a data-dependent mode utilizing a single MS survey scan (*m*/*z* 350–1,650) with a resolution of 60,000, and MS/MS scans of the 15 most intense precursor ions with a resolution of 15,000. The dynamic exclusion time was set to 20 seconds and automatic gain control was set to 3×10^6^ and 1×10^5^ for MS and MS/MS scans, respectively.

### Mass spectrometry data processing

MaxQuant (v2.0.3.0) with implemented Andromeda peptide search engine was used for analyzing the raw MS and MS/MS data. All searches were done on the basis of unspecific protease cleavage, main ion search tolerance of 10 ppm and MSMS tolerance search of 50 ppm and enabling the feature “match between runs”. The reconstituted *in vitro* antigen processing samples were searched against a database containing the sequences of all SARS-CoV-2 proteins (note that Spike and Nucleocapsid were substituted for the sequences of the recombinant ones), cathepsins, MHCII and all reviewed *Spodoptera frugiperda* proteins (Uniprot, access on March 2020) as an internal control. The database used for the cathepsins digestion experiments included only the protein antigen sequence used, and the corresponding sequences of the cathepsins. In both cases a FDR of 0.01 (1 %) was used as well as a decoy database search. All identifications with a FDR higher than 0.01, reverse identifications and contaminants (identified by MaxQuant) were removed for data analysis. Each set of experiments was analyzed together, treating technical and biological replicates as independent samples.

All MS raw files from the reconstituted *in vitro* antigen processing experiments were processed as previously reported in(7). In brief, allotype-specific subset identifications from the evidence file were submitted to the plateau webserver using the same database used for the peptide identification. Each of the consensus peptides identified was then used to determine replication and retrieve a MS1 relative intensity. These relative intensities were averaged throughout the different replicates, and unless otherwise indicated only peptides found in at least 2 technical replicates out of 2 independent experiments were considered. For the proteolytic degradation experiments we took the spectral counts for each peptide identified from the corresponding peptide.txt files. Thus, the number of times that every residue could be mapped to the proteińs sequence.

### Processing of donor samples: PBMC isolation and fractionation

PBMCs isolated from blood samples of either healthy individuals or donors recovered from SARS-CoV-2 infection were isolatrd via density-gradient sedimentation. Thawed PBMCs were first resuspended in complete medium and counting of living cells was performed using trypan blue in a cell counting chamber. When stated PBMCs were used directly in specific experiments, and for others specifically isolated cellular fractions were used. In these cases, cells were isolated by magnetic cell separation using CD14 microbeads for monocytes, and subsequently CD4 microbeads for T helper cells (all Miltenyi Biotec, Bergisch Gladbach, Germany). The homogeneity of the cell preparations was controlled by flow cytometry. For flow cytometric analysis a total of 1×10^5^ PBMCs per well were seeded on 96-well plates, cultured in X-VIVO 15 medium (Lonza, Basel, Switzerland) supplemented with 4 % human AB plasma (Innovative Research, Novi, MI, USA), and provided with the corresponding peptide pools (17.5 ng/ml per peptide).

### ELISpot

Dual secretion of IFNγ and IL-10 was determined using the enzymatic Human IFNγ/IL-10 Double-Color ELISPOT Kit (Cellular Technology, Shaker Hieghts, OH, USA) and using pre-isolated CD14^+^ monocytes and CD4^+^ T cells. 5×10^4^ APC were seeded together with 1×10^5^ CD4^+^ T cells in the presence of relevant peptides at 2.5 ng/ul diluted in X-VIVO 15 medium (Lonza, Basel, Switzerland) supplemented with 4 % human AB plasma (Innovative Research, Novi, MI, USA) in 96-well plates. The cells were then pre-incubated for 48 h, washed, transferred to an ELISpot plate and further incubated for 60 h. The secreted cytokines were determined according to the manufacturer’s instructions, and counting of spots place on an ImmunoSpot analyzer (Cellular Technology). Dual secretion of cytokines was determined by overlapping the corresponding signals (IFNγ-red and IL-10-blue). The extent of the response is directly correlated to the surface area covered by each signal, in this case determined for each colony counted.

### Antibodies and reagents used in cell culture experiments

The following antibodies were purchased from the stated vendors and used according to the manufacturer specifications: for western-blots Rabit anti-6HisTag (Abcam ab1187). In flow cytometry experiments we used: anti-CD4, anti-CD137, anti-CD319 (all Miltenyi Biotec), anti-CD3, anti-PD-1, anti-IL-2, anti-TNFα, anti-IFNγ (all Biolegend). And in case of Elispots we considered: anti-IFNγ (capture and FITC-labelled detection antibodies), anti-IL-10 (capture and biotinylated detection antibodies), anti-FITC HRP (all Cellular Technology).

### Flow cytometry

To accumulate cytokines, cells were treated for 4 h with 5 mg/ml Brefeldin A (Merck, Darmstadt, Germany) after 140 h incubation time. Further, cells were briefly reactivated by addition of 10 ng/ml PMA and 1 mg/ml ionomycin (all Merck) for 1h prior to flow cytometric analysis. Cells were then harvested, Fc-receptors blocked (FcR Blocking reagent, Miltenyi) and stained for extracellular markers (CD3, CD4, CD137, PD-1, and CD319), subsequently fixed with 2 % paraformaldehyde (Morphisto, Offenbach am Main, Germany), permeabilized with 0.5 % saponine (Merck), and stained for intracellular cytokines (IFNγ, IL-2, and TNFα). Samples were acquired on a FACSCanto II flow cytometer with FACS-Diva software (v10, BD Bioscience).

### IEDB data retrieval and analysis

We opted for a scheme, where epitopes flagged as tetramers and with affinity values below a conventional threshold (Aff < 1000nM) were sufficient to define HLA restrictions of a ligand, similar to a recent publication(8). In contrast to this conservative approach, we considered entries eliciting recurrent responses (RF > 0.5, and n>10, named here HF_TcAs here) where the restriction element was unknown. Our DRB1* panel has a decent coverage throughout the individuals in these studies, hence these allotypes are potential restricting elements of those HF_TcAs. Exemplarily, 80 % of the individuals recruited in one of the most comprehensive studies validating immunodominance to SARS-CoV-2 expresses at least one of the allotypes in our DRB1 panel(9). Likewise, if we take into account all epitopes described in TcAs data with associated HLA information, the estimated probability that one of our selected allotypes is the restricting element is higher than 0.5 in approximately 80% of the entries. This value reflects the phenotypic coverage for each individual entry of our panel according to those restrictions stated by these studies.

### Analysis of antigen processing rules and antigenic peptide features

Each protein residue was assigned a value for each of the following parameters: resistance to proteolytic degradation (Res_Prot), relative surface accessible solvent area (rel_sasa) and a pseudo-Sensitivity to endo-protease digestion. For rel_SASA we used the relative values calculated using the pdb PISA tool(10) using as input the structural models available for the Spike and the Nucleocapsid protein as of September 2021 in the Zhang lab website (https://zhanggroup.org//COVID-19/). Res_Prot considers spectral counts for every residue. Pseudo endo-protease cleavage (sensitivity) was retrieved by analyzing proteolytic maps. We deemed residues to be resistant to proteolysis if they met two criteria: i) it was identified in three or more reaction mixtures, and ii) this criterion was met for at least seven consecutive amino acids. Note that this approach yields regions where endo-proteolysis has occurred and at the same time exo-proteases may have trimmed the peptide ends. For this analysis all available entries in the IEDB described as epitopes, ligands or T cell Assay related (TcAs) were equally coded as binary vectors (0: not present, 1: part of a hit) (Accessed Sept 2022). The sum of appearances of each residue as a Hit facilitates defining their relative frequency. If additional data was available, e.g. binding affinity measured experimentally, this information was used to subset those hits. Finally, each candidate epitope identified by the reconstituted antigen processing system was coded at the amino acid level considering the MS1 intensity (e.g. Spike_DRB1*01:01_n for candidate n derived from the Spike protein identified for DRB1*01:01). We worked with either normalized (to the maximum of each vector), count-based of hits or binary-based vectors, as stated in each case.

For the context dependent analysis, we determined the average for each parameter according to the values of each amino acid (Eq. 1). As control we used a randomized pool of peptides excluding all regions labelled as HF_TcAs, Tet_TcAs, MPpN and S.

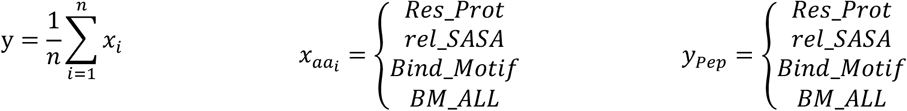

### Statistical analysis and model description

All analysis were carried out using R, version 4.2.1 over the RStudio suite unless otherwise stated. Binary logistic regression models were generated based on the general expression:

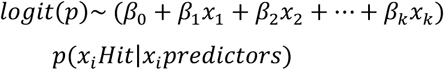

Each predicted candidate from the *in silico* tools (ipEp), identified experimentally (epEp) or its combination (pEp) were considered as single or combined explanatory variables (β_0_ to β_n_) to define a binary response output at the residue level (y = 0 for negative, or y = 1 for positive or selected hits). Different types of IEDB curated entries were defined as response variables following the criteria indicated (Ligand, Tetramers and HF_TcAs).Goodness of fit and the performance of the models were estimated by calculating the Akaike Information Criterion (AIC) and considering the Area Under the Curve (AUC) from Reiceiver Operator Curves (ROC). Likewise, we calculated the positive predictive value of each model as indicated below:

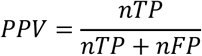

---

**Fig. S1.**
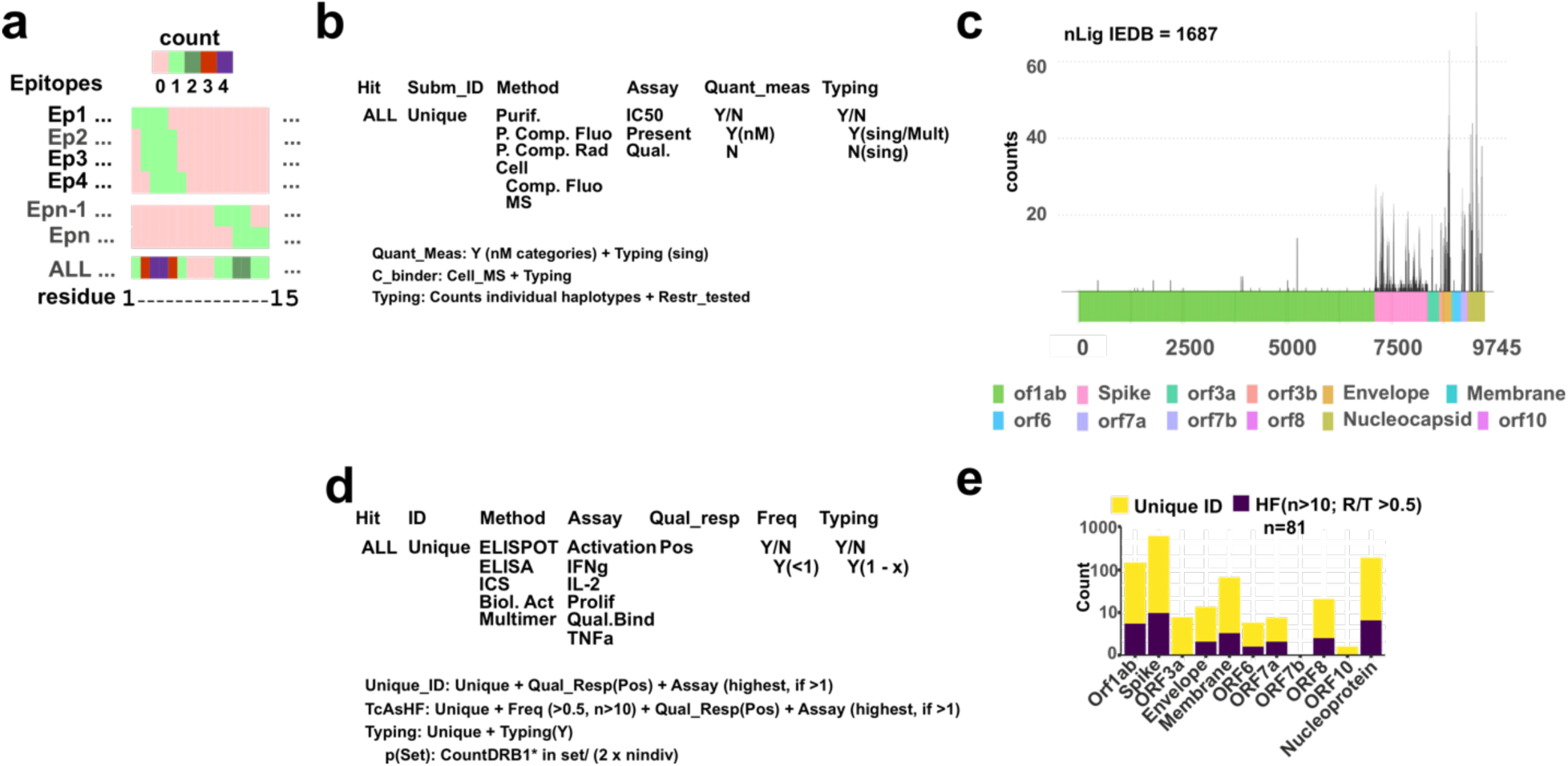
Summary of the retrieval of IEDB data for its use. **a**. Vector coding scheme of all available entries. **b.** Scheme of the data arrangement downloaded and its usage for downstream applications in case of Ligands. There are three subsets of information available and the filtering criteria for each of them is considered. **c**. Schematic representation of the counts of measurements for each residue within the ligand report. **d**. Same as in b) but in this case referred to data derived from T cell assays (TcAs). **e**. Counts of T cell Assay data for each orf considered where the height of the bar represents the total number of unique hits. Those considered as high frequent hits are shown in purple (HF, n individuals tested higher than 10 and at least 5 of them with a positive response).

**Fig. S2.**
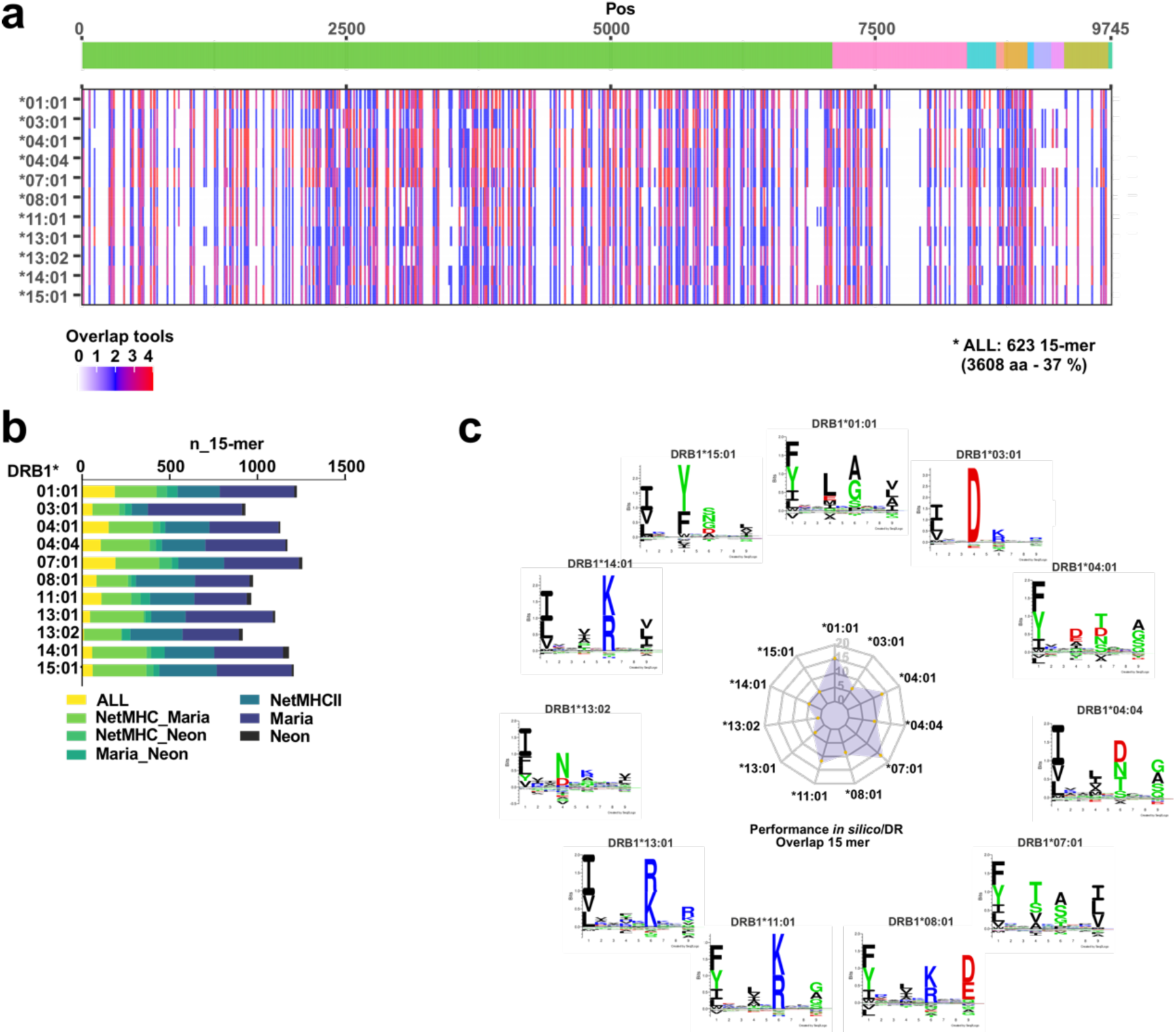
Overview of the performance of the *in silico* tools used to predict binding and/or presentation by the selected set of DR-allotypes. **a.** Summary of the overlap of the different tools plotted over the viral proteome (schematically presented on top) for each allotype selected. Hits are highlighted in colors on a white background (no hit) and the extent of overlap between tools is shown according to the scale shown in the legend. The number of 15-mers (n-15-mer) identified by each tool or combination of tools is shown as a cumulative bar chart. **b.** Global overlap (entire viral proteome) between the different predictors used for each allotype and shown as a bar-chart. **c.** Performance of the *in silico* prediction tools for each allotype, represented as the overlap of 15-mers over the total number of hits per allotype, on the middle radar-plot. On the outer part the binding motifs for each allotype are shown, as reported from eluted data available on NetMHCIIPan4.0.

**Fig. S3.**
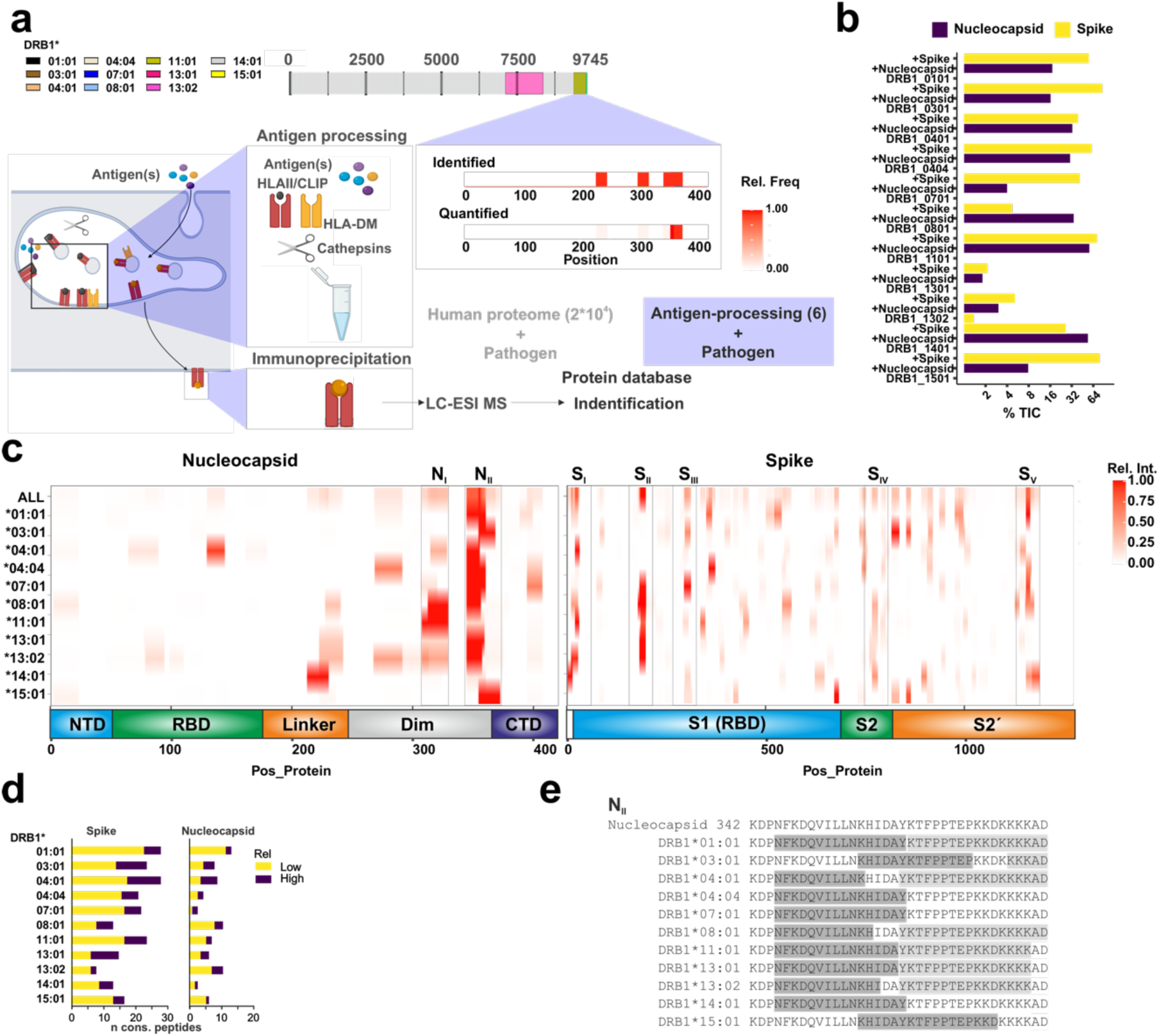
Overview of the reconstituted antigen processing system and its performance. **a**. Rationale and experimental overview of the antigen processing system. The protein database used for the searches consists of 311 entries including common MS-contaminants and abundant proteins from the host used for recombinant protein expression (Exp. Host, *S. frugiperda*), those used for protein manipulation and antigen processing proteins as well as all reference SARS-CoV-2 proteins. **b.** Summary of the performance of the antigen processing system for the selection of candidate antigenic peptides. The sum of the MS1 intensities for the SARS-CoV-2 identified peptides is represented for each of the 3 sets of experimental conditions tested. n = 2 experiments with 3 technical replicates measured. **c**. Summary of the antigenic regions selected by each allotype (stated in the left) over the Nucleocapsid (left) and the Spike (right) protein and depicted according to their relative MS1 intensity. In the lower part the main domains defined for this protein are indicated. The relative frequency of the overlap of candidates selected by the 11 allotypes of our panel is shown in upper row, referred as ALL. **d.** Summary of the number of consensus peptides (maximum of the overlap of the series of nested peptides) identified by MS. The color code refers to the number of peptides defined as those with higher MS1 intensity than the mean (High), or lower (Low). **e.** For each panel, the first line represents the antigenic sequence whereby the first position of the amino acid sequence is provided. Subsequent lines indicate the main (dark background) selected antigenic peptides by each allotype as well as those with lower intensities (gray).

**Fig. S4.**
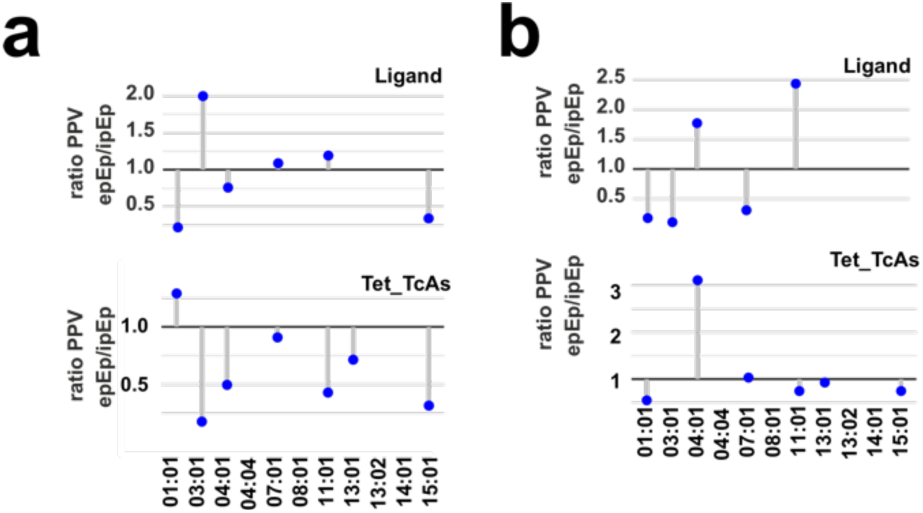
Detailed analysis on the performance of the potential Epitope predictions. **a.** Positive Predictive Value ratios for MS-derived (epEp) and *in silico* (ipEp) models for Ligand and Tetramer identifications for candidates for the **a.** Nucleocapsid, and **b.** Spike protein. A ratio lower than 1 refers to a better PPV of the *in silico* model whereas a ratio higher than 1 refers to a higher PPV of the MS-model for each allotype.

**Fig. S5.**
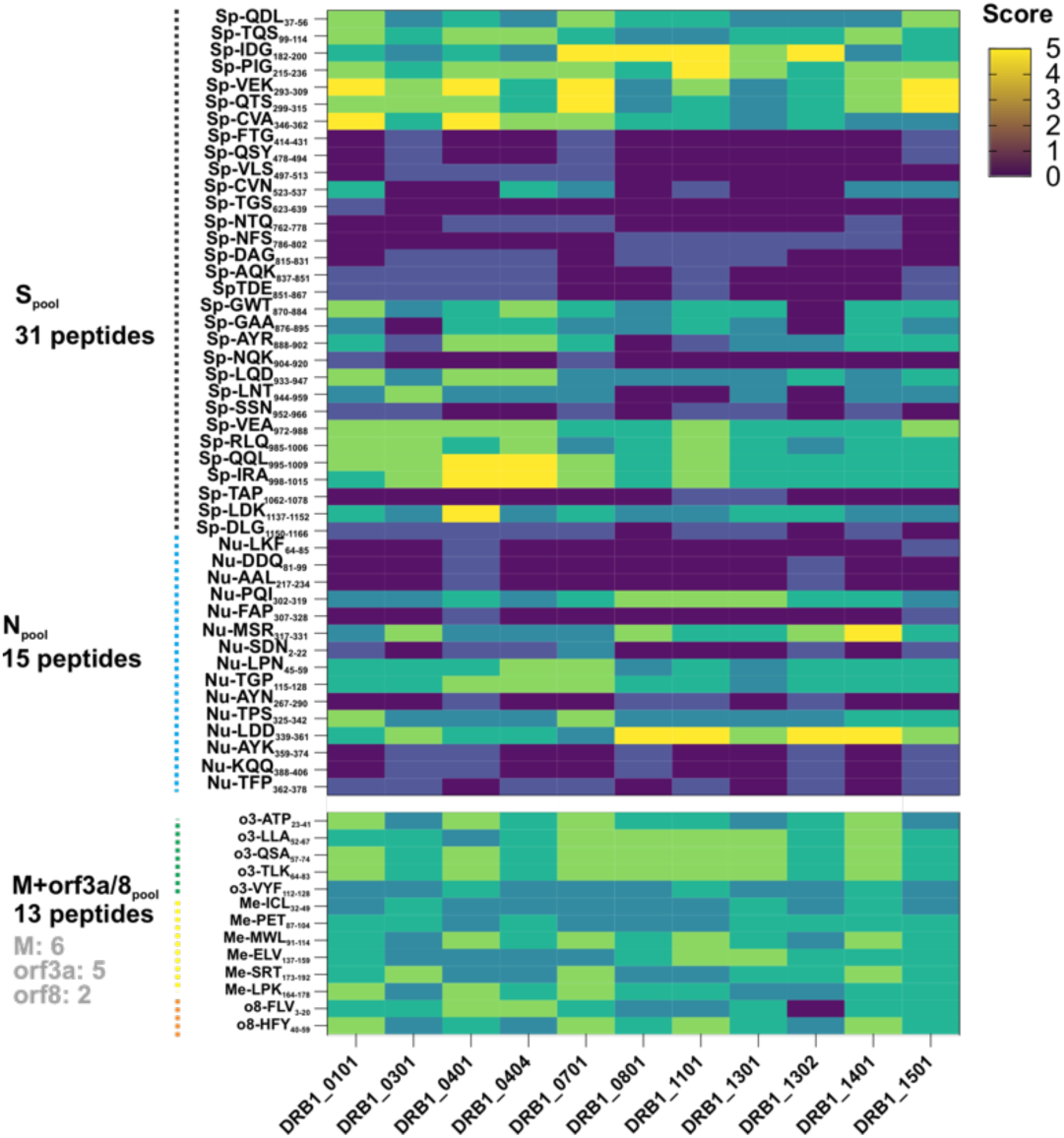
Overview of the minimalistic peptide pools (MPps). Scores indicate the number of times that they were found by any of the predictors used (see scale on the right side). Peptides are pooled by antigens and the total number of peptides of each pool is indicated. For the control peptide pool M+orf3a+orf8 the number of peptides derived from each antigen is indicated in gray. In the names of each individual peptide the name of the antigen and the first amino acid position is indicated.

**Fig. S6.**
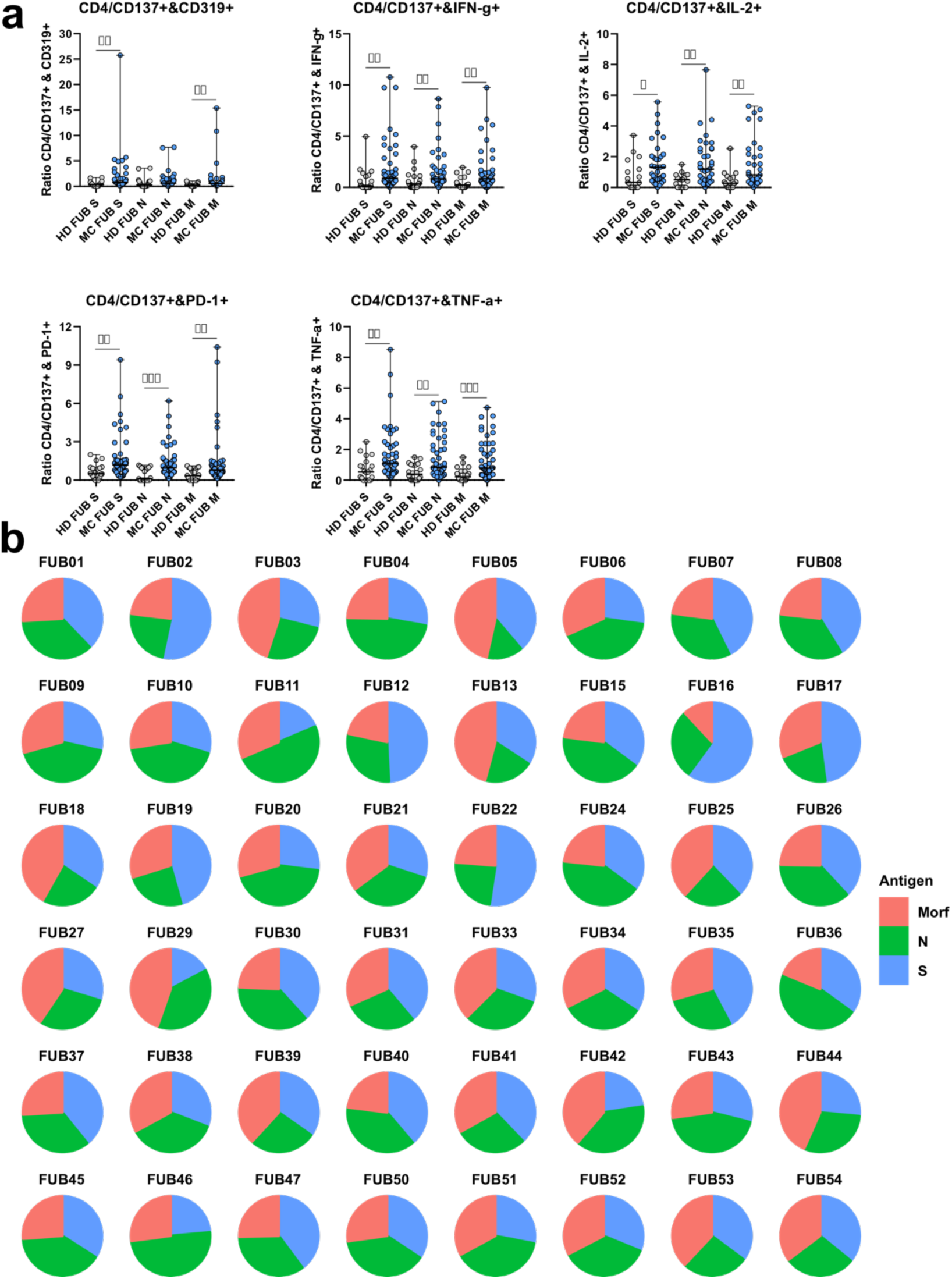
Overview of the activation and profile of T cells responding to the broad coverage peptide panel. **a.** Phenotype of Healthy donor (HD, n=24) and Post-COVID (MC n=48) responder cells to the pools used. Each group was tested for the activation via surface and intracellular cytokine stains upon stimulation with each peptide pool. The difference between the median of the responses was compared applying a non-parametric Mann-Whitney test, and the significance is reported as follows: *p < 0.05; **p < 0.01; ***p < 0.001; ****p < or = 0.0001. **b.** Contribution to the total response of each peptide pool in post-COVID individuals.

**Fig. S7.**
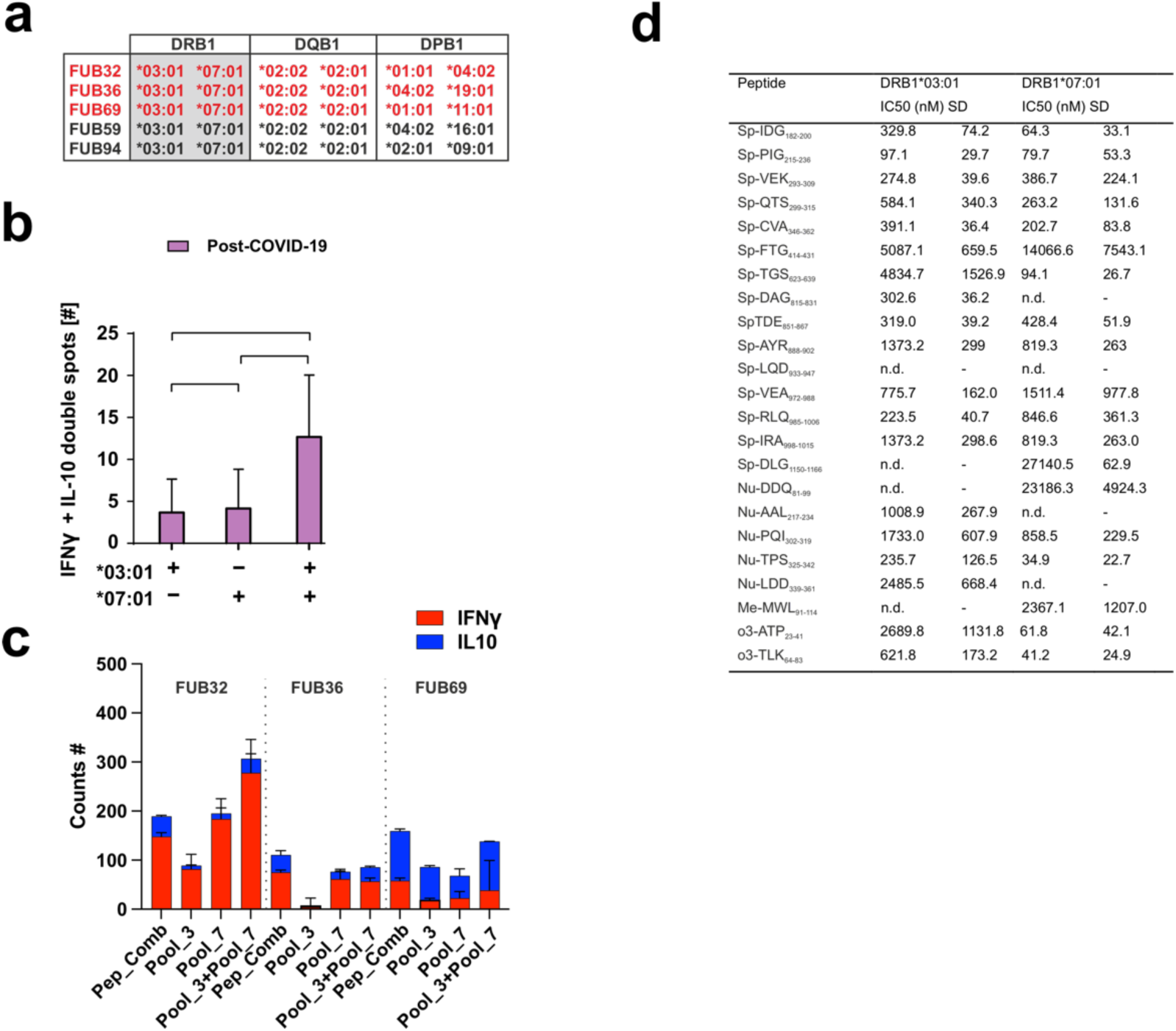
Additional information on the experiments performed with HLA-matched individual’s cells. **a.** Full HLA-types of the donors selected based on the HLA-DRB1* allotypes of interest (grey-shaded boxes). Individuals highlighted in red are those that recovered from SARS-CoV-2 infection. **b.** Number of clones secreting IFNγ and IL-10 for individuals recovered from SARS-CoV-2 infection when primed with peptide pools matching DRB1*03:01, DRB1*07:01 or both. Results show Average and SD of n=2 measurements on 3 individuals. Significance was defined using 1-way Annova with Holm-Sidak correction. * p<0.05, n.s. not significant. **c.** Overview of response achieved for each individual depending on the peptide pool used. Pep_Comb: sum of the responses achieved by the dual peptide combinations; Pool_3: peptide mixture from all those selected as preferential candidates for DRB1*03:01; Pool_7: peptide mixture from all those selected as preferential candidates for DRB1*07:01; Pool_3+Pool_7: the mixture of both peptide pools. The extent of the cytokine response measured for each cytokine is shown colored according to the legend. Measurements represent the average and SD for each individual measured in two technical replicates. **d.** Summary of the binding affinity measurements for the two DRB1*-allotypes. Peptide sequences are indicated as well as the Average IC50 values resulting from n=3 independent experiments measured in triplicates and SD are also indicated. n.d. refers to not determined.

**Figure S8.**
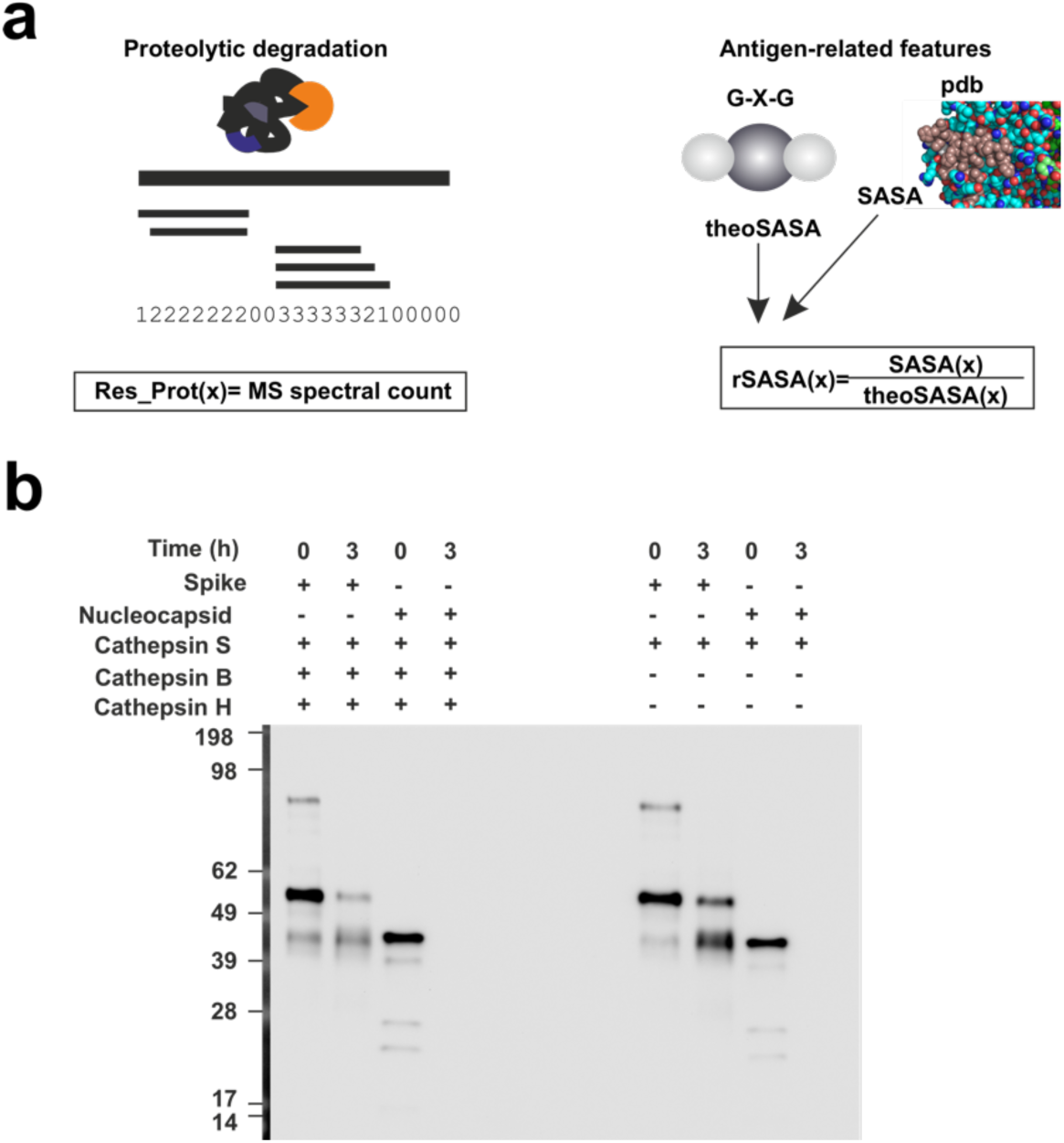
Summary of the main antigen- and antigen-processing related features. **a.** Scheme of the scoring system applied at the residue level for each of the features considered. Proteolytic degradation experiments were performed as indicated and spectral counts are considered to define Res_Prot and Sensitivity to proteases. Structure-related features were calculated from structural models available in the Zhang lab website ((https://zhanggroup.org//COVID-19/) accessed on Sept. 2022 and coded with the corresponding measured values (rel_sasa). **b.** SDS-PAGE demonstrating the *in vitro* degradation of the antigens considered in the presence of the stated proteases for 1.5 and 3h (left), and the mapping of the measured peptides on a scheme of the corresponding antigens.

**Fig. S9.**
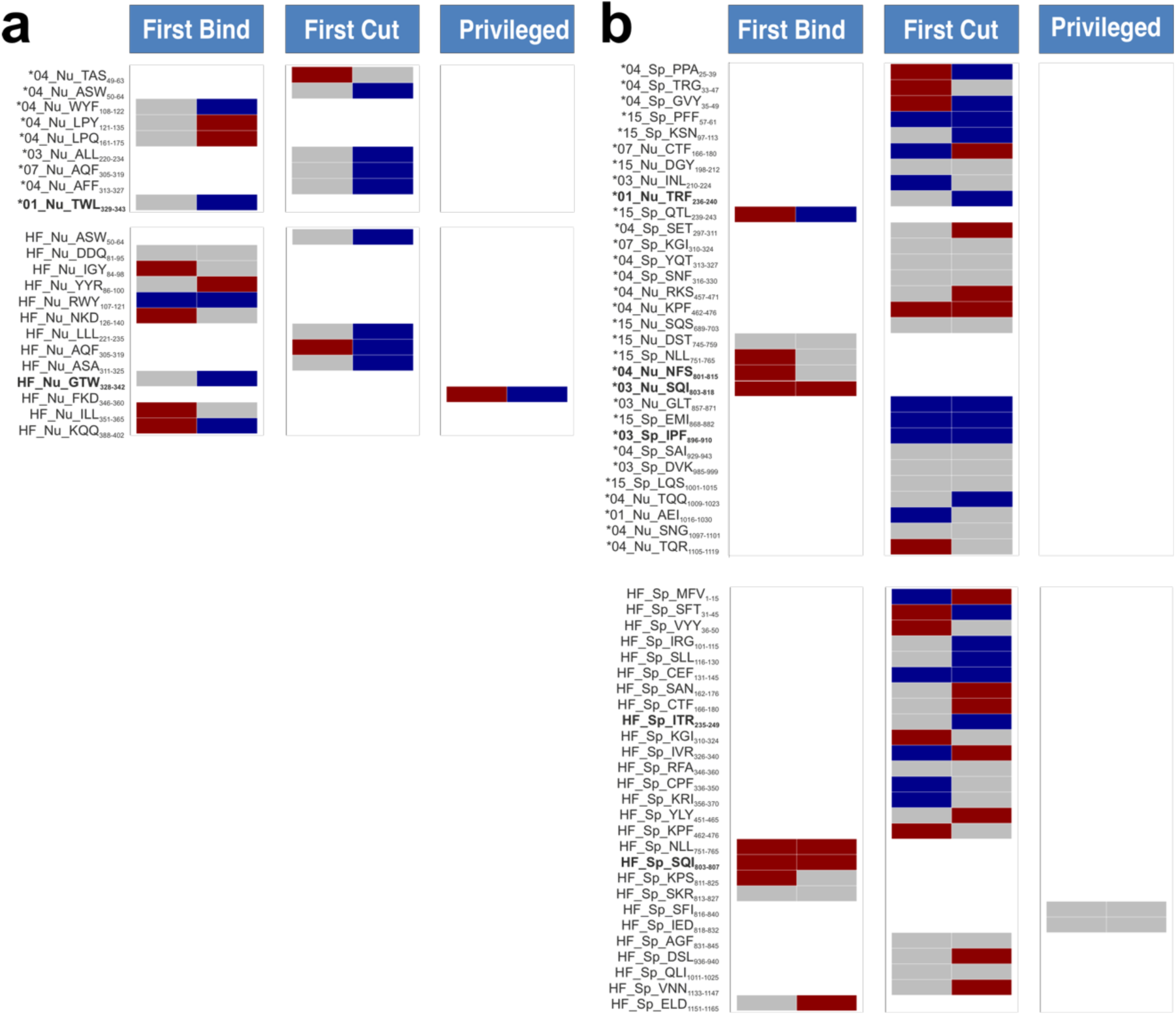
Inferred peptide selection model for all known epitopes from the IEDB. **a**. refers to the Nucleocapsid and **b**. for the Spike. In both cases antigen processing mechanism and antigen-intrinsic features of every known IEDB epitope according to our definition of Tet_TcAs (multimer identified) and HF_TcAs (positive epitopes tested in more than 10 individuals and triggering responses in at least 5) are shown. For Tet_TcAs entries, we indicate the main restriction associated to the peptide at a 2 digit resolution followed but a two characters code for each antigen (Nu for Nucleocapsid and Sp for Spike) and the first three amino acids of the entry followed by the positions covered. Peptides spanning same regions in Tet_TcAs and HF_TcAs and identified with more than one restriction as Tet_TcAs are shown in bold letters. Each model is inferred according to the parameter Sensitivity to proteases, then SASA and Res_Prot averaged values for each entry are considered. Random distributions of averaged feature values for peptides mapping to regions excluding HF_TcAs, Tet_TcAs and MPpN and MPpS are used as controls (note there is one distribution for each antigen consisting of 10 entries for the Nucleocapsid and 20 for the Spike protein). Wilcoxon rank test is used to compare medians of the control distribution with that of the corresponding epitope (significance levels: *p<0.05). Color coding reflects the direction and significance of the deviation from the control median: if the feature value for a given epitope was higher than the median of the corresponding random distribution, a red color was assigned; if it was lower, a blue color was used. Not statistically significant cases are represented in gray.

**Table S1.**
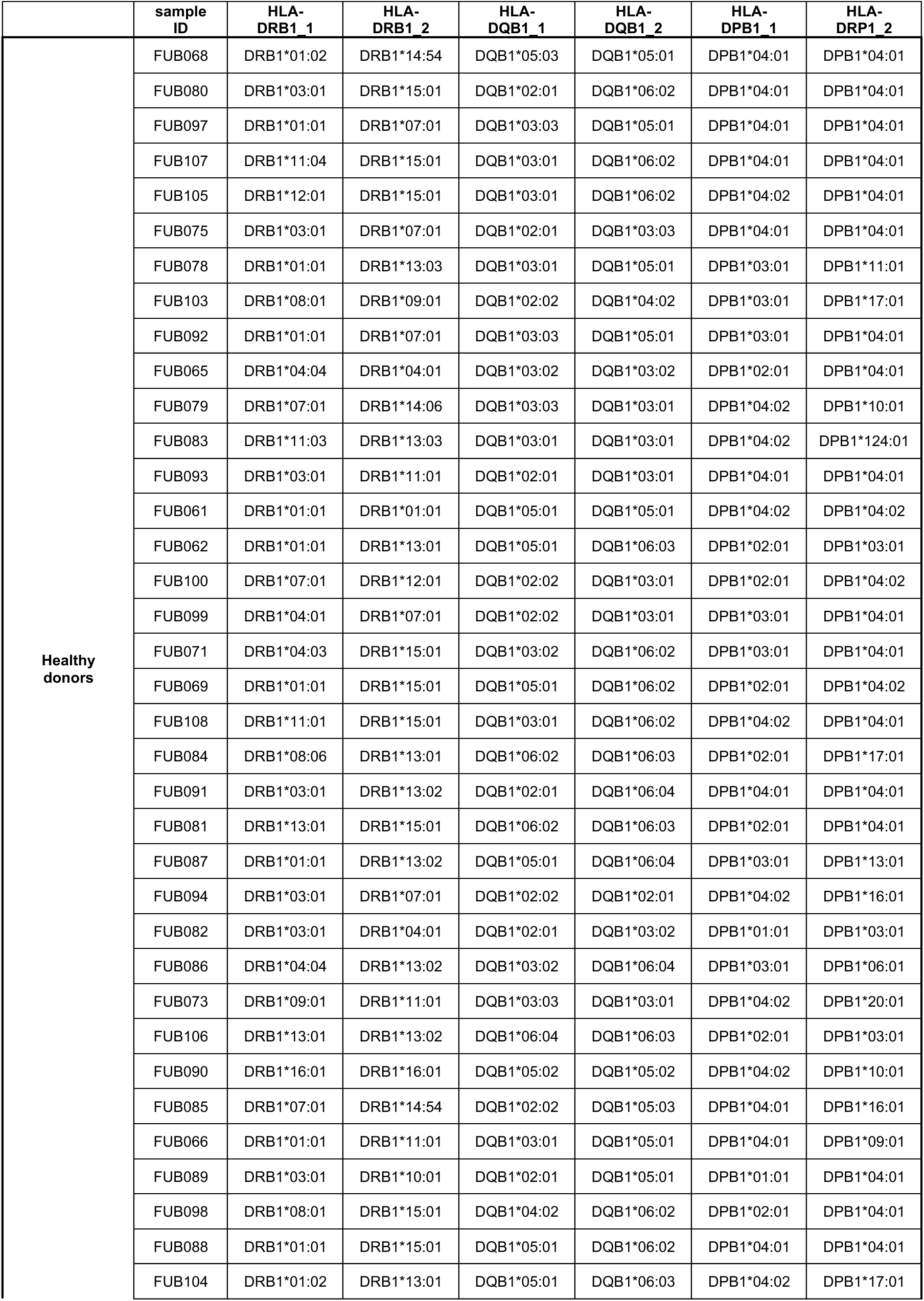
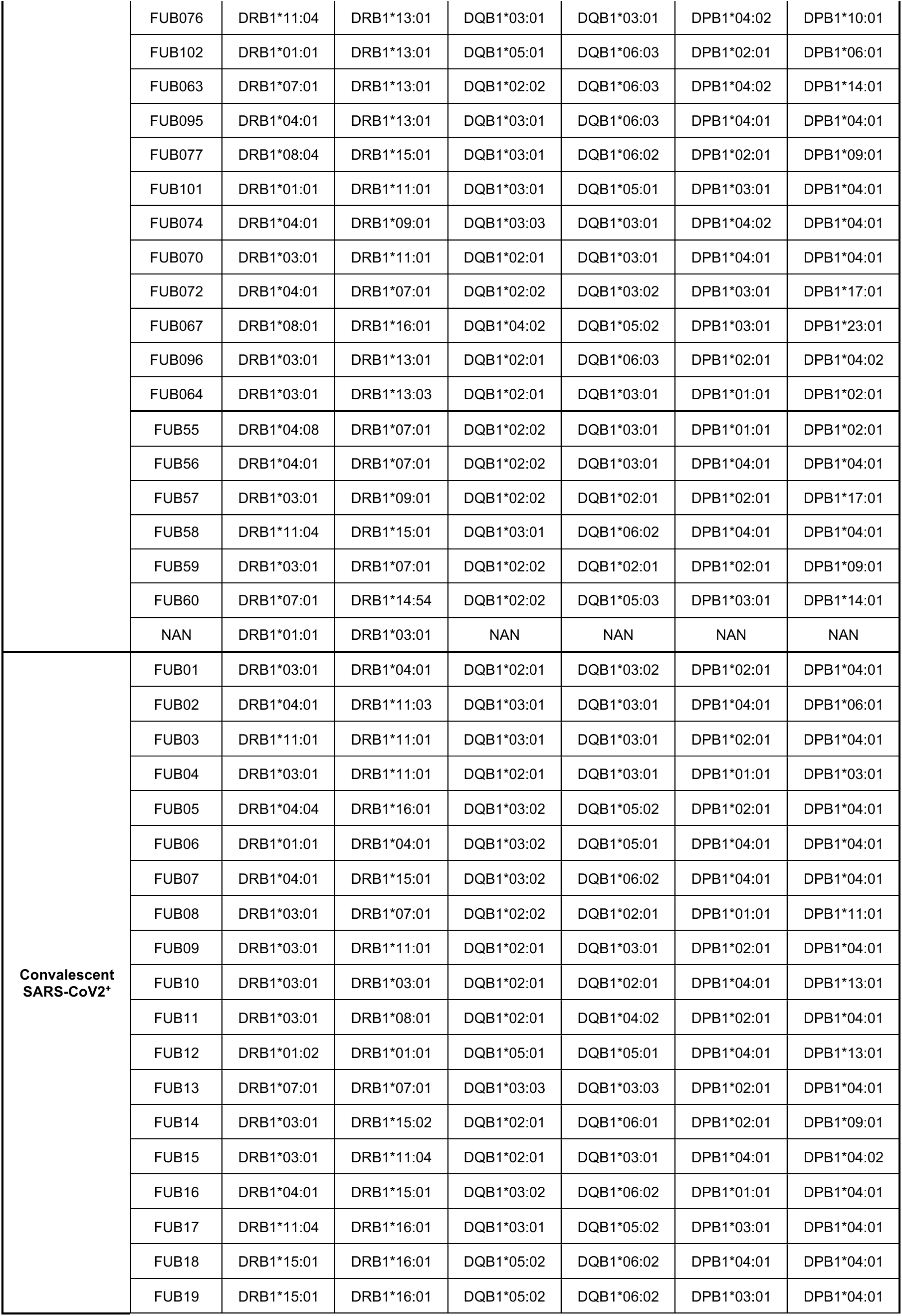
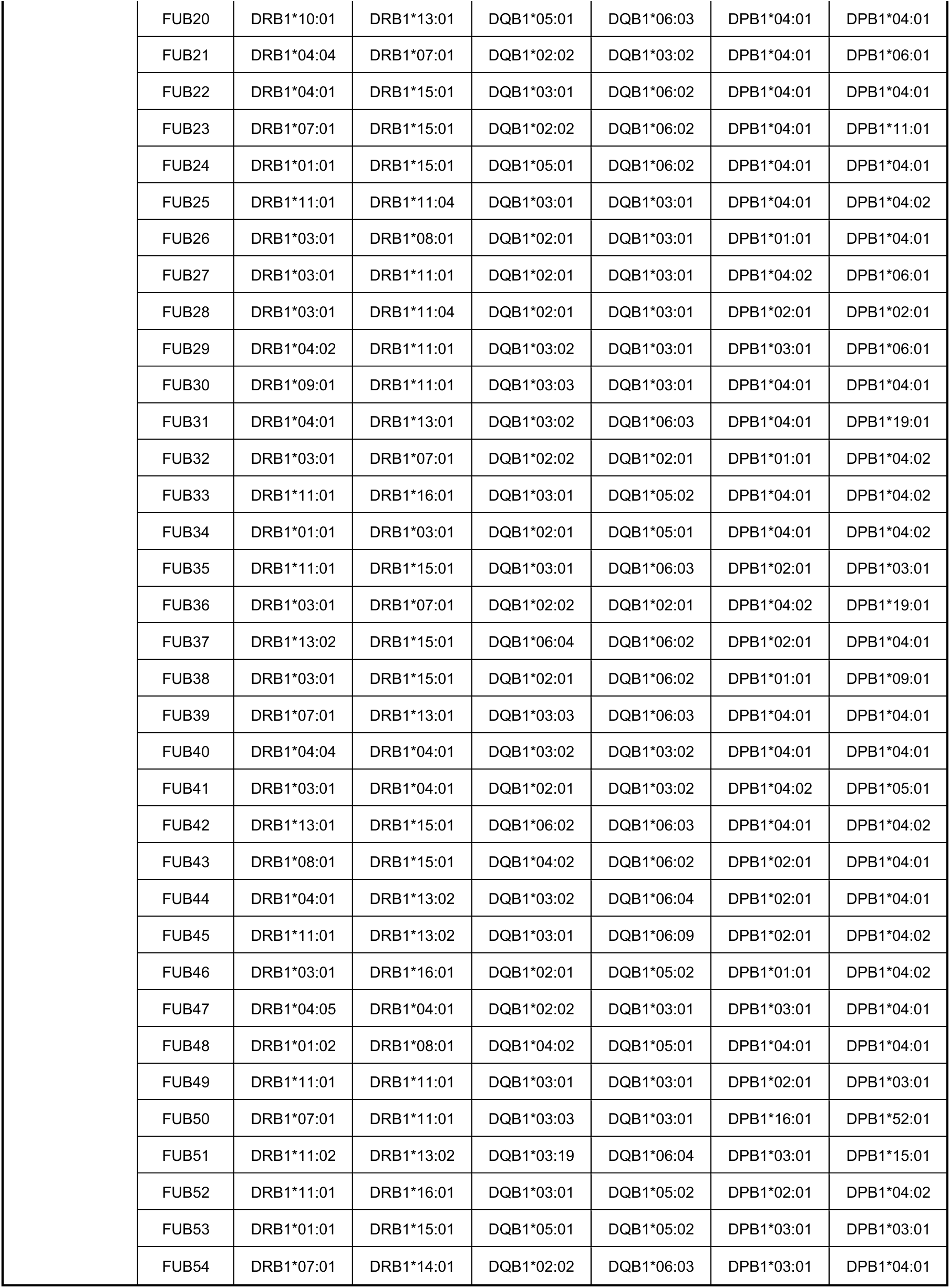
Naming and HLA-type of the 109 individuals recruited for this study.

**Table S2.**
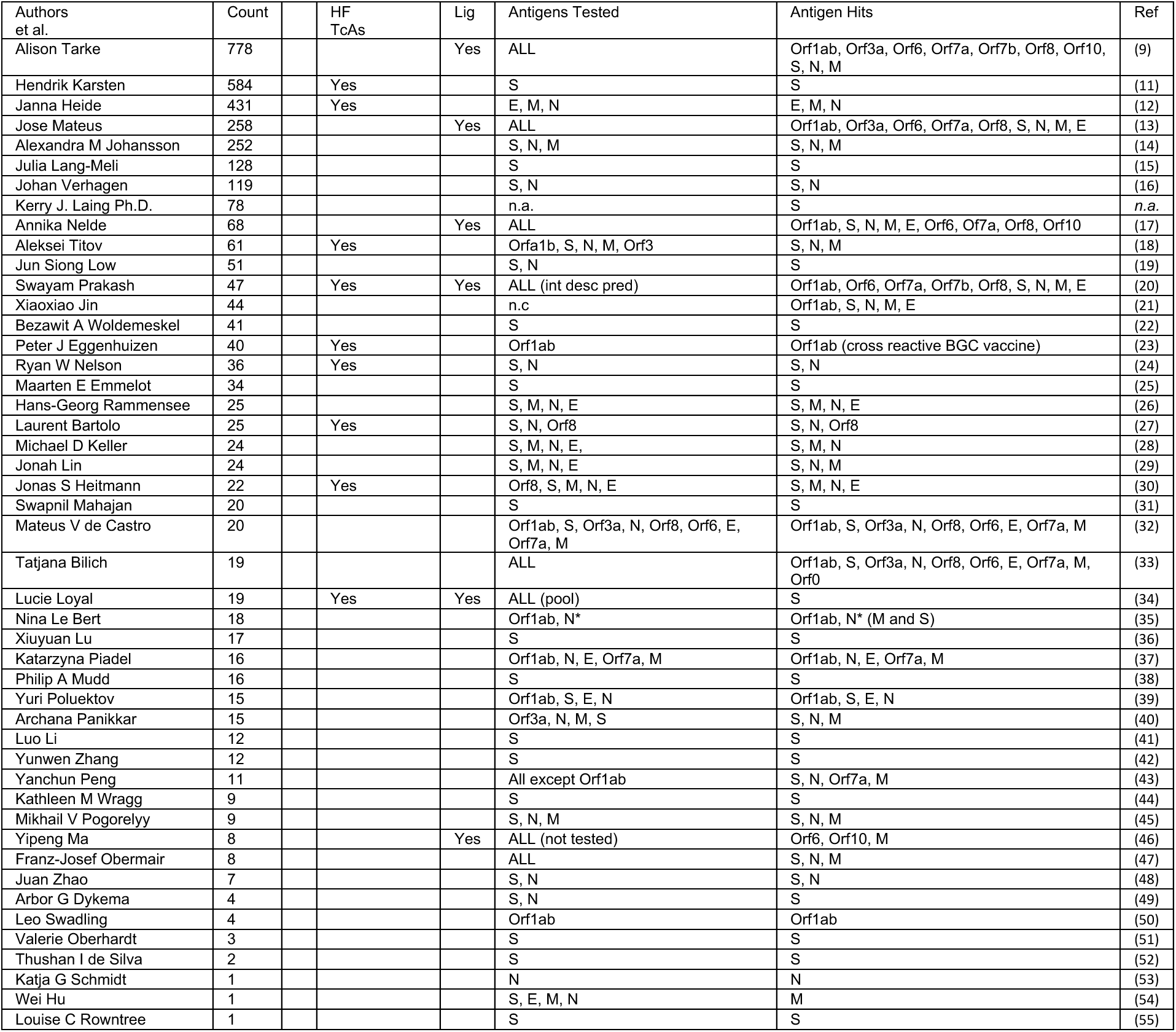
Summary of the T cell assay data retrieved from the IEDB on September 2022. First author of the publication (first column) and entry of the bibliographic reference (last column). “Count” refers to the number of entries yielding positive responses and “HF_TcAs” (Yes) indicate that the study identifies entries that yield positive responses in at least half the tested (more than 10) individuals. “Lig” (Yes) states whether the study compiles ligand information and “Antigen Tested” and “Antigen Hits” refer to the orfs that where considered and yielded responses in the corresponding study.

**Table S3.**
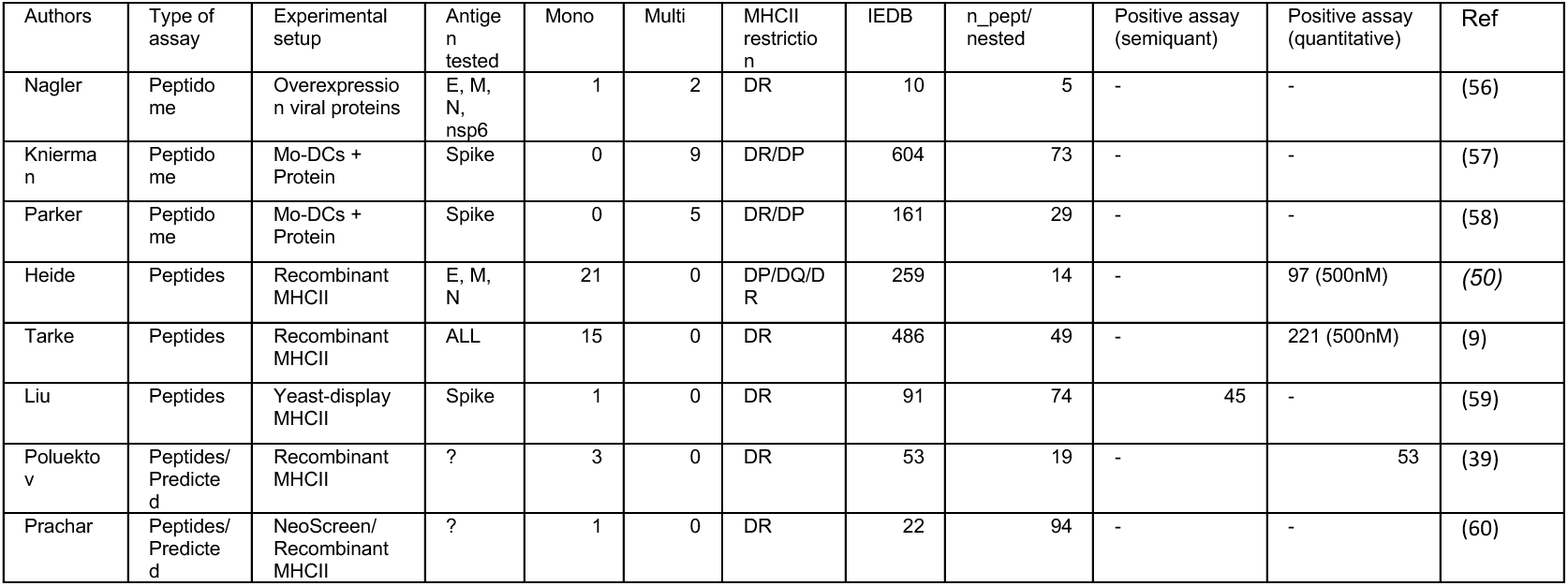
Summary of the Ligand data retrieved from the IEDB on September 2022. First author of the publication (first column) and entry of the bibliographic reference (last column). “Type of Assay” indicate whether the entry was defined upon elution from MHC molecules in an immunopeptidome analysis or using synthetic peptides, and the type of assay that was conducted is referenced in “Experimental setup”. “Antigen Tested” refers to the source(s) of peptides that were considered in the study and “n_peptides/nested” indicates whether the study considers series of nested peptides. “Mono” and “Multi” refer to the background in which ligand binding is assessed, single allotypes in isolation or derived from cellular models in which several allotypes are present. “MHCII restrictions” tested at the molecular level and resulting number of “IEDB” entries. In case of “positive assays”, the number of entries with “semi-“ or “quantitative” information is referred.

**Table S4.**
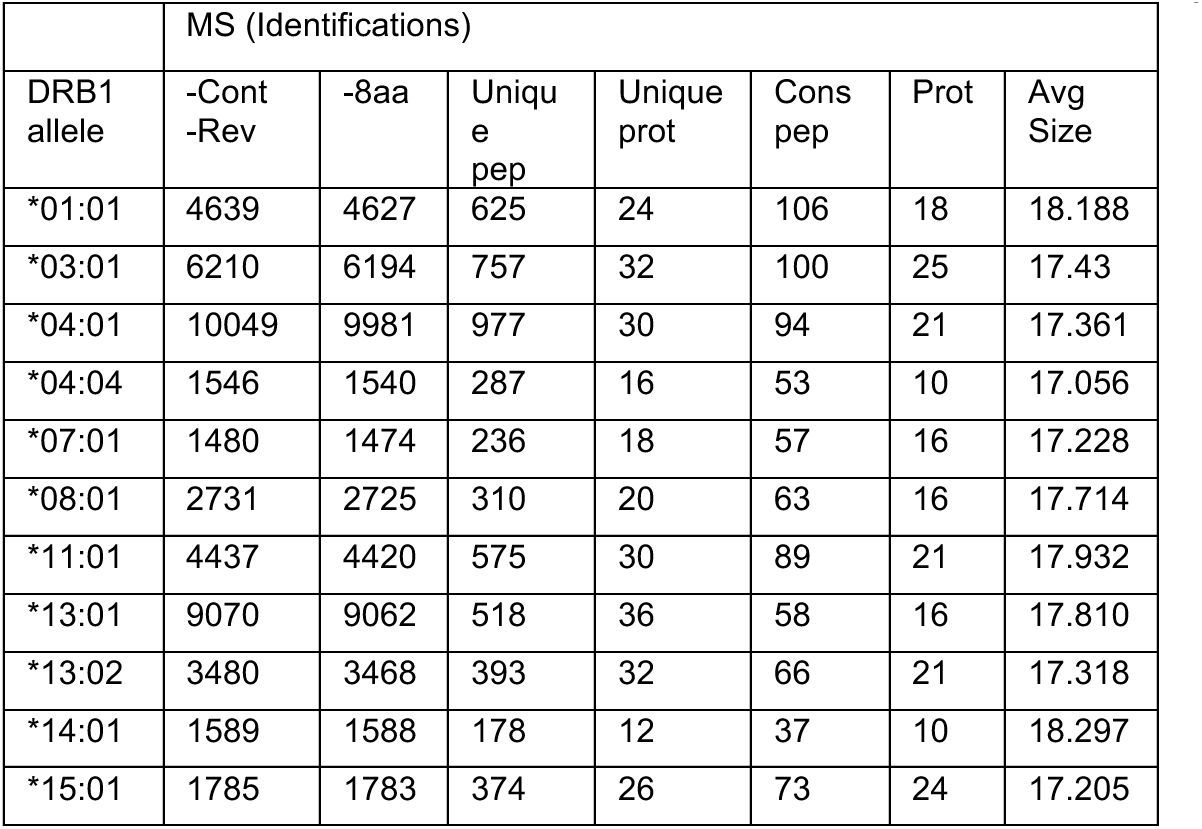
Summary of the *in vitro* reconstituted antigen processing system. Raw files were used to identify peptides present in the samples on MaxQuant using a customized database that included the entire SARS-Cov-2 protein and all recombinant molecules used. All raw files were processed on the same search and the total No. IDs. was 102081, which after filtering contaminants and reverse identification (“-Cont, -Rev”) was reduced to 96689. Peptides of sizes smaller than 8 amino acids were also removed for the analysis (“-8 aa”) and when considering a PEP cutoff of 0.01 was reduced to 47016. “Unique pep” (peptides) or “prot” (proteins) identified are indicated, as well as the number of “Cons pep” (series of nested peptides) and the number of “Prot” (proteins) they derive from. Note as well that the “Avg (average) size” of the eluted peptides is uniformly distributed.

**Table S5.**
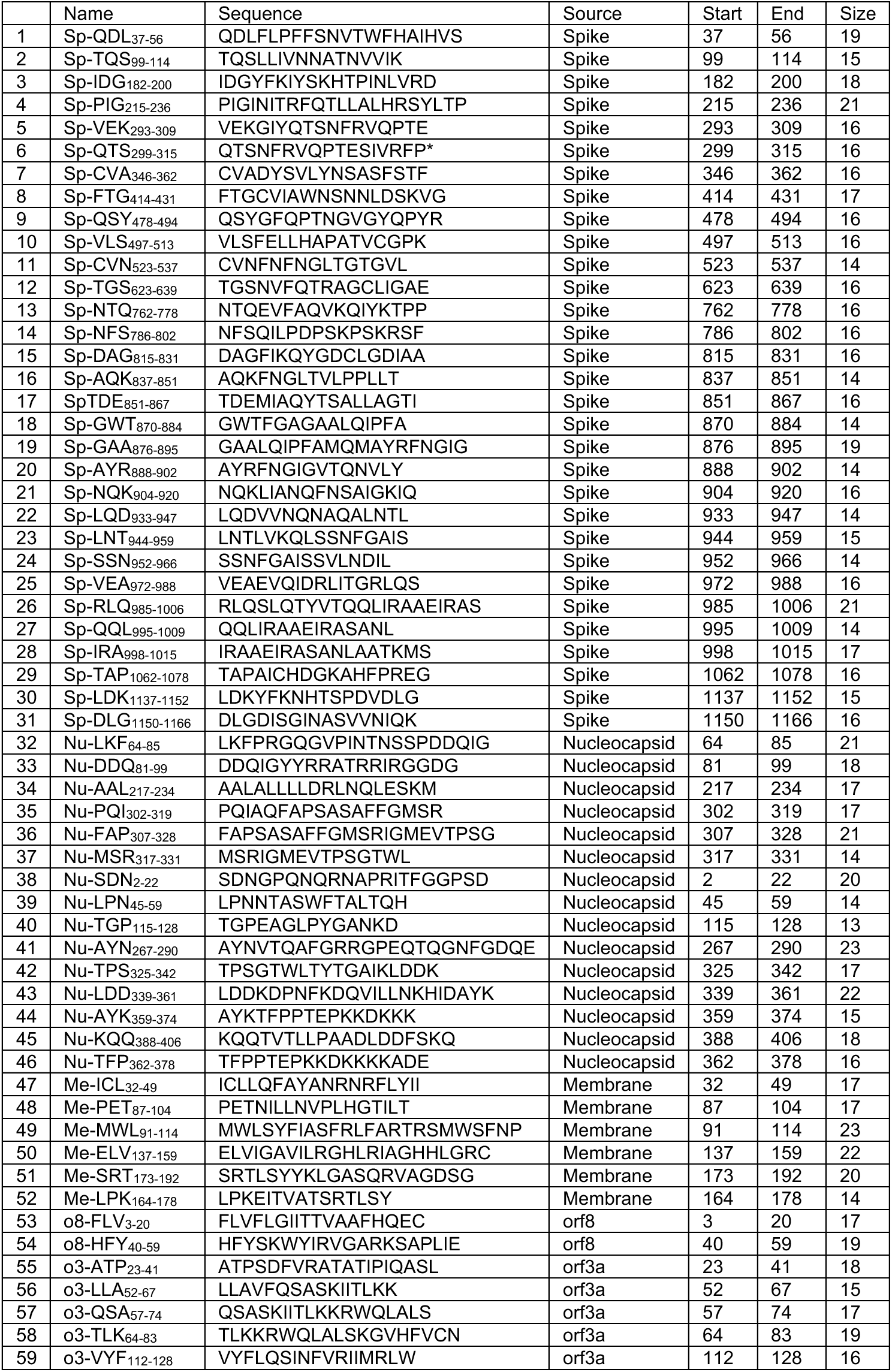
Main features of peptides tested in the different peptide pools used in this study.

